# Transient histone deacetylase inhibition induces cellular memory of gene expression and three-dimensional genome folding

**DOI:** 10.1101/2024.11.21.624660

**Authors:** Flora Paldi, Michael F Szalay, Marco Di Stefano, Daniel Jost, Hadrien Reboul, Giacomo Cavalli

## Abstract

Epigenetic memory enables the stable propagation of gene expression patterns in response to transient developmental and environmental stimuli. Although three-dimensional (3D) organisation is emerging as a key regulator of genome function, it is unknown whether it contributes to cellular memory. Here, we establish that acute perturbation of the epigenome can induce cellular memory of gene expression in mouse embryonic stem cells (mESCs). Specifically, we uncover how a pulse of histone deacetylase inhibition translates to changes in histone acetylation and methylation, as well as global and local genome folding. While most epigenomic and transcriptional changes are readily reversible once the perturbation is removed, genome architecture partially maintains its perturbed conformation. This is significant, as a second transient pulse of hyperacetylation induces continued gene expression deregulation at hundreds of loci. Using ultra-deep Micro-C, we associate memory of gene expression with enhancer-promoter contacts and repressive chromatin topology mediated by Polycomb. These results demonstrate how cells are able to record a transient stress in their 3D genome architecture, enabling them to respond more robustly in a second bout of the same perturbation.

---

Cellular identity is established by gene regulation and epigenetic mechanisms that shape the transcriptional landscape. The information to maintain transcriptional programs is stored in alternative chromatin states that provide the means for cellular plasticity to respond to developmental and environmental cues. This is particularly true for embryonic stem cells (ESCs) that have the developmental potential to give rise to all germ layers. ESCs have distinctive permissive chromatin where activating and repressive configurations often co-exist^1^. These bivalent or poised states are mainly established around developmental genes and require a fine balance between opposing signals to keep gene expression sufficiently low but also prime genes for future activation^2,3^. An important feature of functional chromatin states is the ability to convert short-lived signals to long-lived changes in gene expression – a concept commonly referred to as cellular memory^4^. Cellular memory is widely accepted as an important aspect underlying development and often involves a complex interplay between different epigenetic layers to stabilize gene expression programs following cellular state transitions^5^. However, the crosstalk between epigenetic mechanisms and to what extent can they contribute to memory have been difficult to study due to functional redundancy between components of the epigenetic machinery, as well as the lack of experimental approaches that uncouple gene regulation from cellular memory.

In this study we sought to understand the dynamics of the epigenome during a short-lived disruption of chromatin state balance. To this end, we pulsed mouse ESCs with the histone deacetylase (HDAC) inhibitor trichostatin A (TSA)^6^ that has rapid, global, yet reversible effects on histone acetylation. This latter aspect was critical for allowing us to ask whether such an acute perturbation could trigger cellular memory on the short term. Using a combination of RNA sequencing (RNA-seq), chromatin immunoprecipitation followed by sequencing (ChIP-seq) and ultra-deep Micro-C, we investigate the interplay of gene expression, histone landscape and genome folding at an unprecedented resolution. We place particular emphasis on 3D genome organisation, which is emerging as a key contributor to cellular identity through its role in gene expression control^7^, and we uncover a novel link between 3D genome organisation and cellular memory.

## Results

### A HDAC inhibition pulse leads to global changes in the histone landscape and gene expression

To disrupt the chromatin state balance of mESCs, we pulsed them with TSA for 4 hours (Fig. 1a). To avoid pleiotropic effects, we optimised treatment conditions so that histone hyperacetylation is induced (Extended Data Fig. 1a-b) but the bulk of the acetylome, cell cycle progression and cell viability remain unchanged (Extended Data Fig. 1c-d). Calibrated ChIP-seq indicated that acute TSA treatment efficiently induced genome-wide H3K27 hyperacetylation (Fig. 1b) resulting in significant increase in H3K27ac signal at >30,000 sites (Extended Data Fig. 1e). We also detected ∼2,800 previously strong H3K27ac peaks that showed modestly decreased but dispersed signal enrichment upon TSA treatment. While H3K27ac loss was highly specific to TSSs, H3K27ac gain occurred ubiquitously around *cis*-regulatory elements, along gene bodies as well as in intergenic regions (Fig. 1c, Extended Data Fig. 1f).

**Figure 1.**
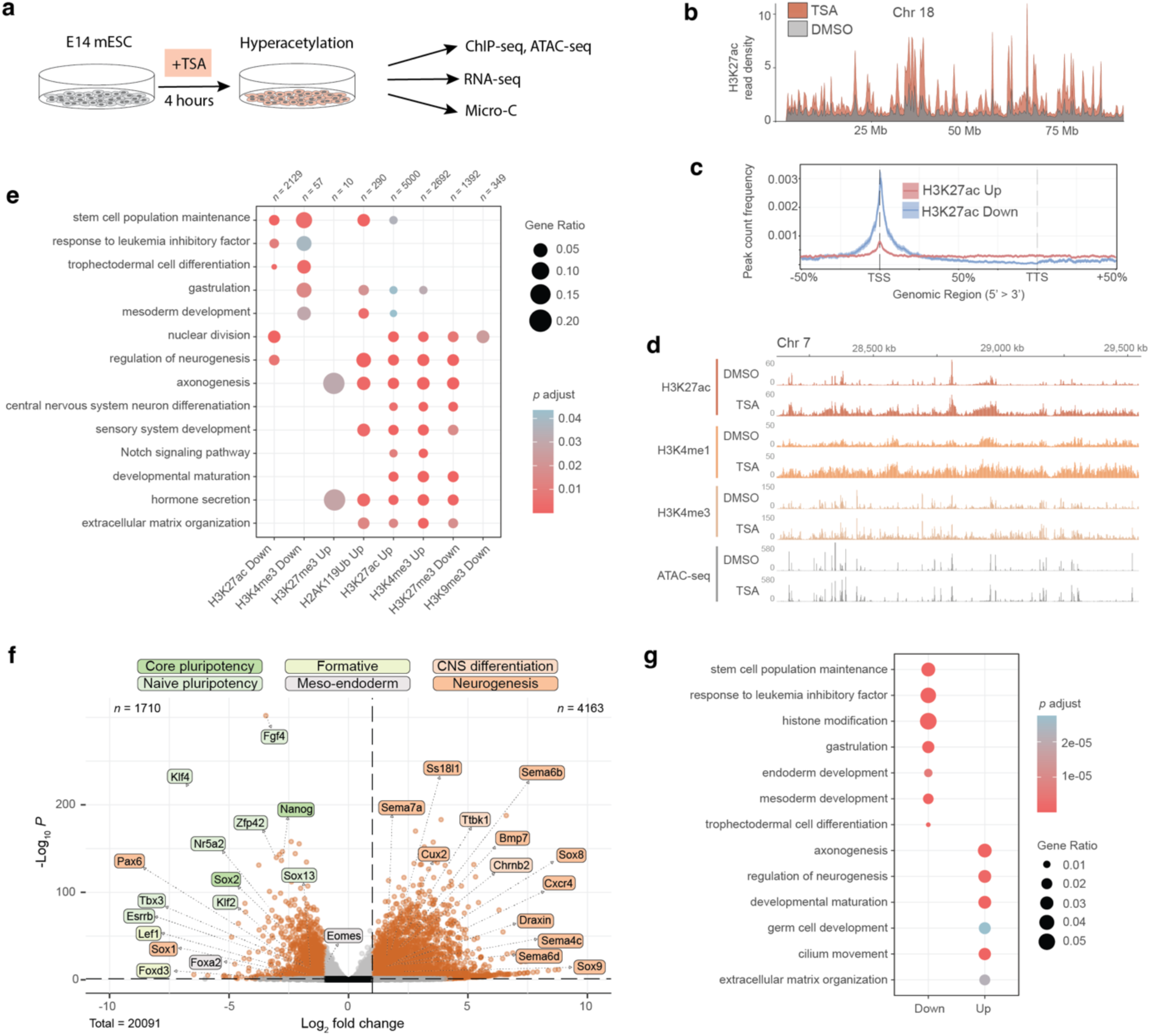
Acute HDAC inhibition leads to global changes in the histone landscape and gene expression. **a**, E14 mESCs were pulsed for 4 hours with Trichostatin A (TSA) and assayed for changes in chromatin states by a combination of ChIP-seq, ATAC-seq, RNA-seq and Micro-C. **b**, Plot showing H3K27ac ChIP-seq read density on chromosome 18 (bin size = 10 kb). **c**, Differential H3K27ac peak count frequency distribution relative to gene bodies (TSS = transcription start site, TTS = transcription termination site). Shading represents 95% confidence interval (CI). **d**, Genomic snapshot of spike-in normalised ChIP-seq and ATAC-seq signal over a typical hyperacetylated region. **e**, Gene Ontology enrichment of terms related to development among genes located within 1 kb distance from differential ChIP-seq peaks. **f**, Volcano plot showing differential gene expression (significance cutoffs: adjusted *p*-value > 0.05, absolute log_2_ fold change > 1) upon TSA treatment. Labels correspond to core and naïve pluripotency, formative, meso-endodermal, central nervous system (CNS) differentiation and neurogenesis marker genes. **g**, Gene Ontology enrichment of terms related to development among up- and down-regulated genes.

As functional chromatin states are maintained by an interplay between active and repressive histone modifications^5,8^, we next focused on characterising secondary changes in the histone landscape. By categorising genomic intervals as active (enriched in H3K27ac, H4K4me1 or H3K4me3) and repressive (enriched in H3K9me3, H3K27me3 or H2AK119Ub), we found that TSA treatment caused a larger fraction of the genome to be in an active state, while repressive intervals decreased nearly 2-fold (Extended Data Fig. 1g). Indeed, we identified thousands of differential peaks for all examined histone modifications, where activating marks were in general gaining signal and repressive marks underwent mostly loss of enrichment (Fig. 1d, Extended Data Fig. 1f). Gene annotation of differential ChIP-seq peaks pointed to an amplification of developmental processes and a suppression of pluripotency (Fig. 1e).

Next, we carried out bulk RNA-seq to discern whether the effects of HDAC inhibition on the transcriptome reflect the changes we detect in the histone landscape. As HDAC1 is an important regulator of early development^9,10^, we focused on transcriptional changes related to ESC identity. As suggested by reorganisation of the histone modification landscape, downregulated genes were associated with stem cell population maintenance as well as endoderm and mesoderm development (Fig. 1f-g). In parallel, we observed upregulation of genes associated with developmental maturation, in particular specification of the neural lineage as previously described^10–13^.

In sum, we found that acute HDAC inhibition leads to widespread accumulation of H3K27ac and global changes in the histone modification landscape that promote a gene expression programme associated with exit from pluripotency.

### Global and fine-scale architectural changes characterise the TSA chromatin state

Imaging studies have shown that TSA treatment leads to chromatin decompaction both at global^14,15^ and local^15,16^ scales. To understand how these alterations translate to changes in chromatin contacts, we generated ultra-deep Micro-C contact maps with 8.5 and 6.6 billion unique valid 3D chromatin contacts for control (DMSO) and TSA conditions respectively (Supplementary Table 1). Our datasets show more uniform genomic coverage over both active and inactive genomic intervals with respect to previous Micro-C studies^17,18^, and provide unbiased genome-wide interaction maps with unprecedented detail (Extended Data Fig. 2a-b). Unlike recent high-resolution capture studies, we do not observe microcompartments at previously described loci^19^, only mild on-diagonal insulation at microcompartment anchor sites. We attribute the differences in our dataset to differences in Micro-C protocol rather than to insufficient data depth (Extended Data Fig. 2c-d).

When we compared contact maps generated from control and TSA-treated cells, we detected a dramatic increase in *trans* contacts upon TSA treatment (Extended Data Fig. 3a). This was concomitant with a marked decrease in *cis* interactions at nearly all genomic distances (Extended Data Fig. 3b). TSA treatment also led to a loss of prominent A compartment interactions - a characteristic of embryonic stem cells^20^ - without major changes in compartment identity (Fig. 2a, Extended Data Fig. 3c). While *trans* contacts were gained in both compartments (Extended Data Fig. 3d), BB interactions - a feature of differentiated cells^20^ - became prominent in *cis*. This observation prompted us to examine if A and B compartments were asymmetrically impacted by TSA treatment. Indeed, gain in activating histone marks in A compartment exceeded that in B, and most gene expression deregulation corresponded to TSSs located in the A compartment (Fig. 2b-c, Extended Data Fig. 3e).

**Figure 2.**
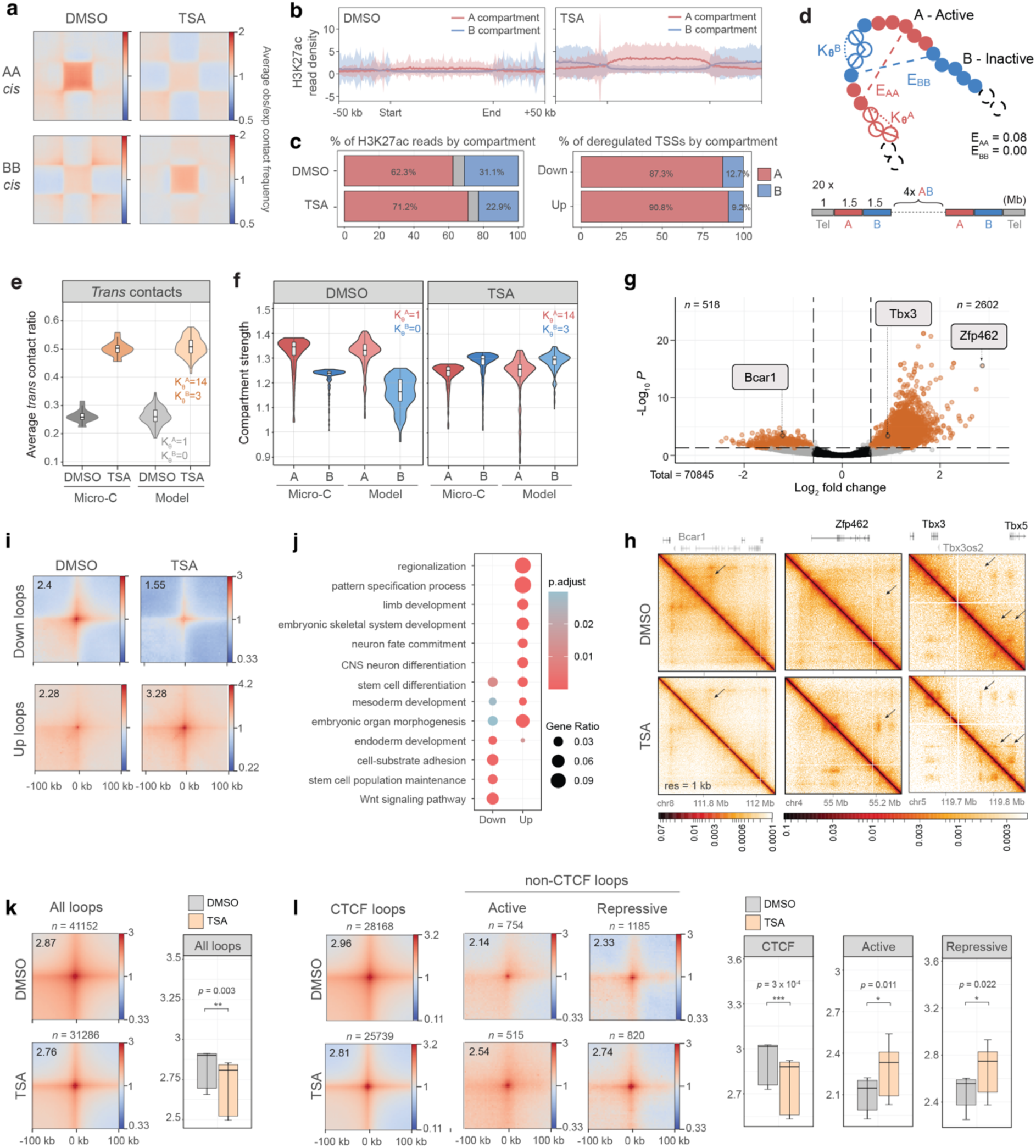
Global and fine-scale architectural changes characterise the TSA chromatin state. **a**, Aggregate plots of homotypic interactions between and A and B compartments in *cis*. **b**, Metaplots showing H3K27ac normalised ChIP-seq read density over A and B compartments (bin size = 1 kb). Shading represents standard deviation. **c**, Distribution of H3K27ac ChIP-seq reads (left panel) and up- and down-regulated TSSs (right panel) by compartment. **d**, Schematic representation of biophysical modelling where E_AA_ and E_BB_ correspond to the attraction energies and K_θ_^A^ and K_θ_^B^ correspond to the stiffness of the chromatin fibre, in A and B domains, respectively. Each chromosome was modelled by a 20 Mb chain where each bead represents 5 kb of DNA. A and B domains were set at 1.5Mb in size to match the mean compartment size derived from the Micro-C data, resulting in 6 A-domains, 6 B-domains, and 2 telomeric regions of 1 Mb each at the extremities for the chain. A nucleus was modelled using 20 chains. **e**, *Trans* contact ratio in DMSO and TSA in the Micro-C data and in the model. **f**, Compartment strength in DMSO (left panel) and TSA (right panel) in the Micro-C data and in the model. **g**, Volcano plot of differential loops between DMSO and TSA (significance cutoffs: adjusted *p*-value > 0.05, absolute fold change > 1.5). Positive log_2_ fold change indicates stronger interaction in TSA. **h**, Contact maps showing differential looping at the Bcar1, Zfp462 and Tbx3 loci. **i**, Aggregate plots of Micro-C signal around differential loops at 4 kb resolution. **j**, Gene Ontology enrichment among genes closest to differential loop anchors. **k**, Pile-ups showing Micro-C signal around all loops identified in DMSO (top) and TSA (bottom) (resolution = 4 kb). Quantification of aggregate loop signal is shown on the right (paired two-tailed t-test; **p < 0.01). Data shown are the median, with hinges corresponding to interquartile range (IQR) and whiskers extending to the lowest and highest values within 1.5× IQR. **l,** Pile-ups of Micro-C signal around loops stratified by the presence (CTCF loops) or absence (non-CTCF loops) of CTCF ChIP-seq peaks at loop bases (resolution = 4 kb). Non-CTCF loops have been further divided to active and repressive based on the presence of activating (H3K27ac, H3K4me1) or repressive (H3K27me3, H3K9me3) ChIP-seq signal at loop bases. Quantification of aggregate loop strength is shown on the right (paired two-tailed t-test; *p < 0.05, ***p < 0.001). Data shown are the median, with hinges corresponding to IQR and whiskers extending to the lowest and highest values within 1.5× IQR.

It has been previously suggested that elevated transcriptional activity or acetylation level increases the stiffness of the chromatin fibre, leading to an increase in *trans* interactions^21^ and chromosome reorganisation^22^. We tested this idea by simulating TSA-induced changes on global chromatin folding using mechanistic 3D polymer modelling^23^. First, we used a single-chain block copolymer model (Fig. 2d) made of A and B chromatin regions to infer self-attraction energies (E_AA_, E_BB_) that best reproduce the A/B compartment strength in DMSO condition (Extended Data Fig. 3f-g, Methods). Since A and B compartments are asymmetrically^22^ affected by TSA-induced hyperacetylation and transcriptional upregulation (Fig. 2b, c), we then simulated the effect of TSA treatment by changing the stiffness differentially within A and B domains in a multi-chain system. By optimising the value of stiffness to match the median *trans*-interaction ratio (Fig. 2e, Extended Data Fig. 3h-j), the TSA conformation was compatible with an increase of stiffness in A-domains to a greater degree than in B-domains (*K*_*θA*_ = 14 *K*_*θ*_, versus *K*_*θB*_ = 3.0 *K*_θ_). Interestingly, the change in stiffness we applied to reproduce the *trans*-contact ratio automatically predicted the swap in compartment strength without modifying the A-A and B-B attraction energies (Fig. 2f, Extended Data Fig. 3i): A-domains lost compartment strength while B-domains gained it, so that in TSA B-B interactions were stronger than A-A. Additionally, our model predicted a displacement of A-domains towards the periphery of the simulated spherical nucleus^24^ (Extended Data Fig. 3k).

Subsequently, we sought to comprehensively map conformational changes that occur at the submegabase scale. Whole genome analyses identified >3000 focal interactions with differential looping strength (Fig. 2g-i) that were strongly associated with developmentally important loci (Fig. 2j, Extended Data Fig. 2l). As we detected nearly 3500 CTCF peaks that became stronger in TSA, we tested whether increase in loop strength was due to increased CTCF binding (Extended Data Fig. 2m). Contrarily, we found that CTCF-mediated loops globally became weaker. Instead, non-CTCF loops – carrying either active (H3K27ac, H3K4me1, H3K4me3) or repressive (H3K9me3, H3K27me3, H2AK119Ub) chromatin signatures – became stronger upon TSA treatment (Fig. 2k-l).

Taken together, TSA treatment had a profound effect on global genome folding, promoting inter-chromosomal contacts and decreasing A-A compartment interactions. These changes in global chromatin folding were compatible with a compartment-specific increase of stiffness of the chromatin fibre in our biophysical modelling simulations. In parallel, TSA treatment caused specific, fine-scale restructuring where CTCF-dependent and epigenetic-state driven loops behaved differently. Importantly, annotation of sites with rewired chromatin contacts mirrors ongoing developmental processes that we identified from changes in the transcriptome and the histone landscape.

### Changes in histone modification landscape and chromatin looping underlie differential gene expression

We next asked what epigenomic changes underlie TSA-induced gene expression deregulation. As upregulated TSSs were enriched for H3K27ac peaks that gain signal in TSA, whereas peaks that showed decreased signal were more frequently localised to downregulated TSSs (Fig. 3a), we conclude that transcriptomic changes are least partially correlated to changes in the H3K27ac landscape. Analyses of epigenetic signatures around upregulated TSSs revealed that gain in H3K27ac was concomitant with a modest increase in chromatin accessibility and a substantial gain of H3K4me1 in surrounding regions. (Fig. 3b) On the other hand, we detected overlapping H3K4me3 and H3K27me3 signal at a large subset of upregulated genes, indicating that bivalent genes are susceptible to TSA-mediated gene derepression. This occurred with a gain in H3K4me3 and H3K27ac, but without a detectable loss of H3K27me3 signal (Fig. 3c, Extended Data Fig. 4a).

**Figure 3.**
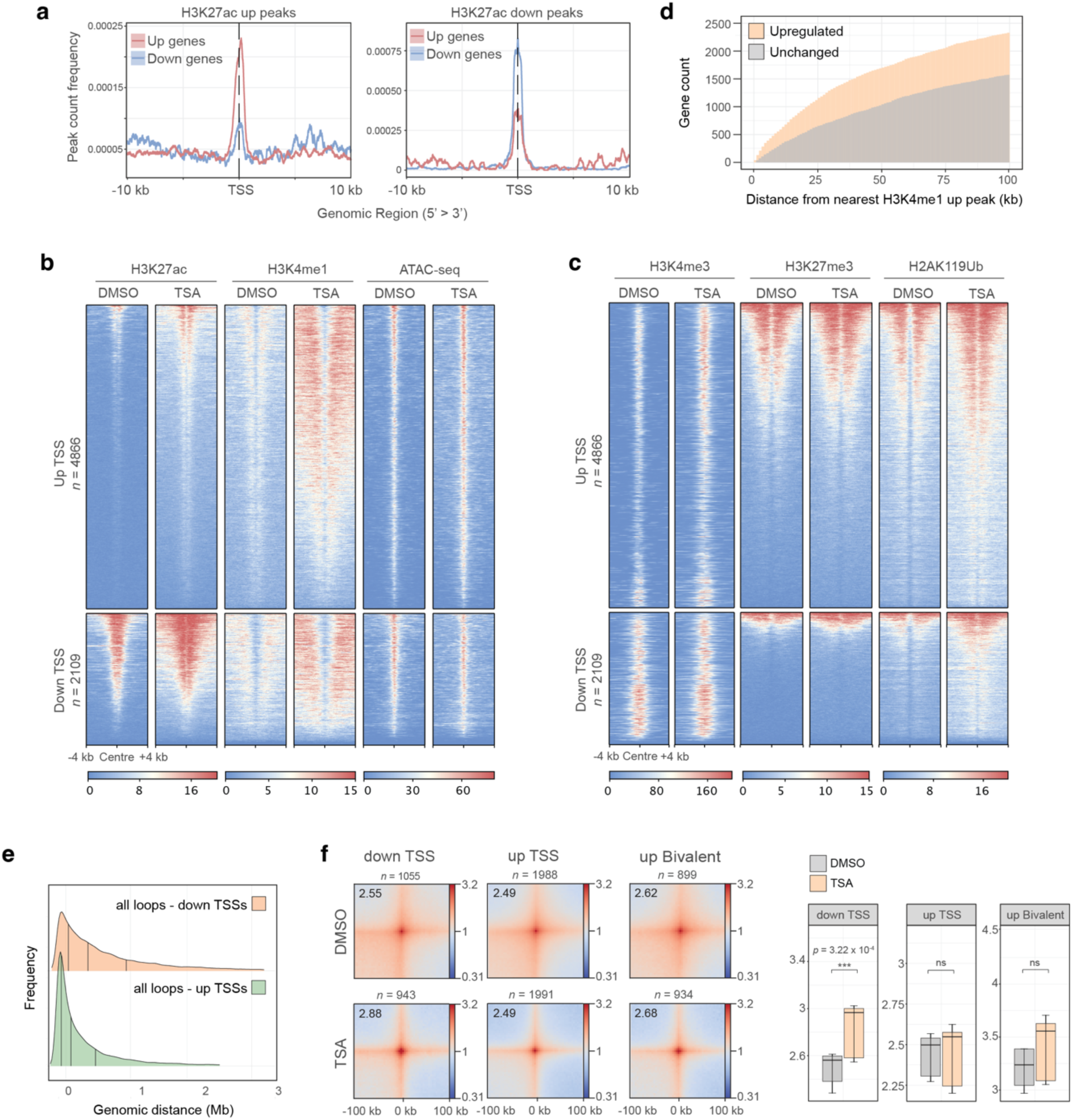
Changes in histone landscape and chromatin looping underlie differential gene expression. **a**, H3K27ac up (left panel) and down (right panel) peak count frequency distribution relative to TSSs of up- and down-regulated genes. Shading represents 95% CI. **b**, H3K27ac, H3K4me1 and ATAC-seq signal around transcription start sites (TSS) of differentially expressed genes. **c**, Heatmaps showing H3K4me3, H3K27me3 and H2AK119Ub signal around TSSs of differentially expressed genes. **d**, Cumulative histogram showing genomic distance between upregulated gene promoters and the nearest increased H3K4me1 peak. Control genes represent an expression-matched gene set that does not increase in expression. **e**, Ridge plot showing the frequency of loop anchors in the function of genomic distance from the nearest deregulated TSS. **f**, Aggregate plots of Micro-C signal around loops where anchors overlap with down- (left) or up-regulated (middle) TSSs, as well as bivalent (right) TSSs that undergo upregulation (resolution = 4 kb). Quantification of piled-up loop signal is shown on the right (paired two-tailed t-test; ns > 0.05, ***p < 0.001). Data shown are the median, with hinges corresponding to IQR and whiskers extending to the lowest and highest values within 1.5× IQR.

To understand if transcriptional upregulation could be caused by ectopic enhancer activation, we examined the linear proximity of genomic regions that gain H3K4me1 signal to gene promoters. This revealed that such peaks form closer to upregulated gene promoters than promoters of expression-matched control genes that do not undergo upregulation (Fig. 3d). Similarly, upregulated TSSs were found to be in closer proximity to previously described primed (H3K4me1+, H3K27ac-) and poised (H3K4me1+, H3K27me3+, H3K27ac-) enhancers^25,26^ (Extended Data Fig. 4b). As enhancer activation and subsequent gene expression are often accompanied by the establishment of enhancer-promoter (E-P) contacts^20,27,28^, we analysed changes in chromatin looping associated with upregulated TSSs. Although we detected an increased linear proximity of loops to upregulated TSSs, gene expression upregulation seemingly occurred without noticeable changes in promoter contacts (Fig. 3e-f).

Interestingly, we observed similar changes in the histone modification landscape at downregulated TSSs, where active marks were gained in the absence of accumulation of repressive histone modifications (Fig. 3a, d). Instead, even without TSA treatment a part of downregulated TSSs were found to be strongly enriched for Myc and YY1 binding (Extended Data Fig. 4c), transcriptional regulators that are direct targets of HDACs and whose acetylation state has been described to modulate their molecular function^29,30^. This suggests that TSA-induced gene downregulation may be partially due to effects on non-histone targets of HDACs. Crucially however, unlike upregulated TSSs, promoter loops around downregulated TSSs became stronger (Fig. 3f). We noticed that around some of the most strongly downregulated genes prominent *de novo* loops formed in TSA-treated cells. In these cases, looping was globally associated with gene downregulation and occurred between existing H3K9me3 sites with only a mild increase in H3K9me3 level (Extended Data Fig. 4d-h). This suggests that repressive chromatin contacts could provide means for gene downregulation even when activating histone modifications are acquired.

Altogether, we find that gene upregulation as well as downregulation occur with a gain of activating chromatin marks. In general, while gene upregulation is potentially associated with enhancer over-activation without gain in E-P contacts, downregulation is linked to repressive chromatin looping.

### ESCs recover their transcriptional identity and chromatin states following the removal of HDAC inhibition

Next, we asked if cells could revert to their unperturbed state upon TSA removal. We postulated that if chromatin perturbation had strictly instructive roles, then the TSA cellular state should be readily reversed once the perturbation is removed. Conversely, if the perturbed chromatin state is a carrier of cellular memory, the changes should outlast the initial causative event. To this end, we washed TSA-treated cells and let them recover for 24 hours, which corresponds to approximately two cell doubling times. After 24 hours of TSA removal, original H3K27ac levels and the rest of the acetylome appeared restored at the protein level (Fig. 4a, Extended Data Fig. 1a-b), indicating successful removal of the perturbation. The effects of TSA on the histone landscape were likewise readily reversible. Excess H3K27ac, H3K4me1 and chromatin accessibility were restored at once (Fig. 4b) and H3K27ac peaks re-gained their enrichment around TSSs (Fig. 4c). Consistent with restoration of chromatin marks, TSA-induced global transcriptional deregulation was nearly completely reversed with only a handful of genes (*n* = 164) showing sustained changes (Fig. 4d). Even at these loci, transcription was largely reversed, and only residual dysregulation persisted. Although we detected increased H3K27ac level at certain genomic loci, these loci were not correlated with genes that remained deregulated (Fig. 4e). Altogether, these data show that histone marks and gene expression are almost entirely restored upon removal of TSA, showing little or no memory of the past perturbation.

**Figure 4.**
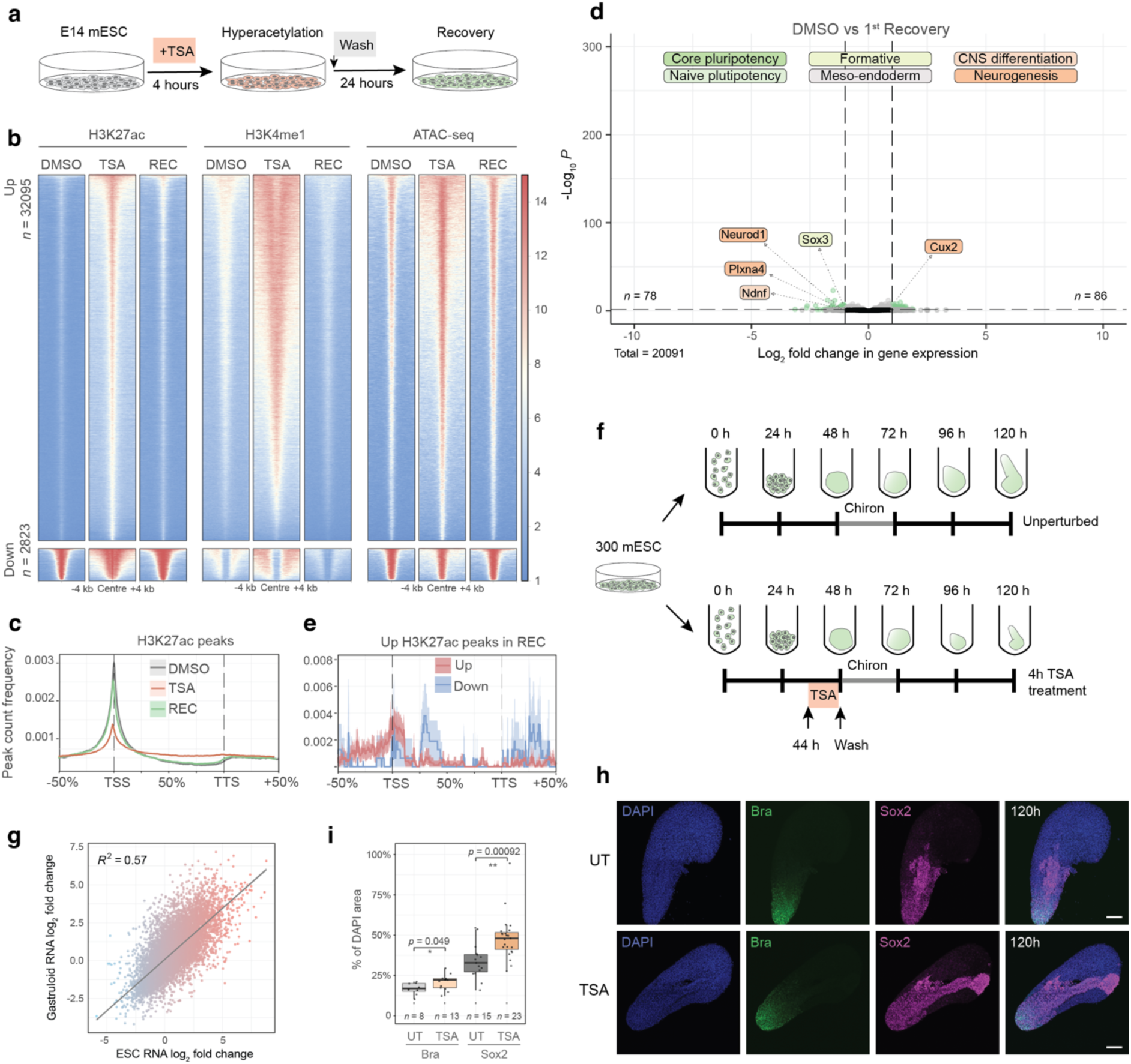
Pluripotent cells recover their transcriptional identity and chromatin states following the removal of HDAC inhibition. **a**, mESCs were washed extensively and were re-assayed for their chromatin states 24 hours later. **b**, Heatmaps H3K27ac, H3K4me1 and ATAC-seq signal in DMSO, TSA and recovery (REC) at differential H3K27ac peaks identified in TSA. **c**, H3K27ac peak count frequency distribution in DMSO, TSA and recovery datasets relative to gene bodies of all genes (TSS = transcription start site, TTS = transcription termination site). Shading represents 95% CI. **d**, Volcano plot showing differential gene expression 24 hours after TSA washout. Labels show developmental marker genes that remain above the significance threshold (adjusted *p*-value > 0.05, absolute log_2_ fold change > 1). **e**, H3K27ac peak count frequency distribution in recovery dataset relative to gene bodies that remain up- and down-regulated (TSS = transcription start site, TTS = transcription termination site). Shading represents 95% CI. **f**, Schematics of TSA treatment between 44 and 48 hours and subsequent washout during gastruloid differentiation. **g**, Scatterplot showing the correlation of TSA-induced transcriptomic changes in mESCs and in 48-hour gastruloids. Shading represents log_2_ fold change in mESCs. **h**, Representative images of Sox2 and Brachyury immunostaining in 120-hour untreated gastruloids (top) or gastruloids 3 days following transient TSA treatment (bottom) (scale bar = 100 *μ*m). **i**, Quantification of Brachyury (Bra) and Sox2 area over DAPI in 120 h gastruloids with and without TSA treatment. Data shown are the median, with hinges corresponding to IQR and whiskers extending to the lowest and highest values within 1.5× IQR (unpaired two-tailed t-test; *p < 0.05, ***p < 0.001).

Importantly, the activity of the pluripotency network was efficiently recovered, and developmental processes were downregulated anew. As mESC culture conditions are designed to actively suppress differentiation, we were wondering if this efficient recovery was a general feature of pluripotent cells, or whether it was due to culturing conditions that impose such a dominant cell state that they overpower the effects of perturbations. To test this idea, we grew mESCs into gastruloids^31,32^, that we treated with TSA for 4 hours immediately prior to the chiron pulse (Fig. 4f, Extended Data Fig. 5a, b). The effects of TSA on the transcriptome strongly correlated between mESCs and gastruloids (Fig. 4g) with GO enrichment analysis showing similar developmental deregulation in both conditions (Extended Data Fig. 5c).

Following washes - which restored H3K27ac within 24 hours (Extended Data Fig. 5c,d) - we let control and TSA-treated cells develop for three days, until reaching a mature gastruloid state. Gastruloids were then assayed for their transcriptomic and morphological features. Interestingly, while the transcriptome was largely re-established, immunofluorescence staining revealed morphological aberrations in TSA-treated (Extended Data Fig. 5e) gastruloids. Although maintaining their ability to specify germ layers, we found that the TSA pulse led to a marked expansion of the area staining positive for the neuroectodermal marker Sox2, while the distribution of cells expressing the meso-endodermal marker Brachyury remained unchanged (Fig. 4h-i).

In sum, mESC possess a remarkable capacity to recover their transcriptional and histone modification landscape following a hyperacetylation pulse. Our findings in gastruloids fall in line with these observations, namely that the effect HDAC inhibition on the transcriptome is profound, but transcriptional recovery from it is nearly complete. Nevertheless, the incomplete recovery of the Sox2 expression domain in gastruloids might indicate that a memory of the TSA pulse might be ingrained.

### Genome architecture retains partial memory of the past conformation

To further explore whether the TSA pulse could be recorded by mESCs, we analysed 3D genome folding upon restoration of the initial conditions. Surprisingly, we found that chromatin conformation did not fully recover: the *cis-trans* ratio was restored only partially (Extended Data Fig. 6a) and contact decay curves showed that *cis* contact depletion persisted, particularly in the A compartment (Extended Data Fig. 6b). In terms of compartment interactions, while AA interactions were efficiently recovered in *trans* and showed some increase in *cis*, BB in *cis* interactions remained prominent following recovery from TSA (Fig. 5a). We could equally detect sustained changes in genome conformation at the gene level. The depth of our data allowed us to carry out eigenvector decomposition at 4 kb resolution^33^. This revealed local instances where transcriptional and architectural recovery became uncoupled. For example, gene expression upregulation at the F11, Klkb1 and Cyp4v3 loci in TSA moved the ∼200 kb long encoding genomic segment to the A compartment. Following recovery, expression of all three genes was successfully restored, however the encoding genomic segment had a continued A compartment identity, like in the TSA condition (Fig. 5b). Additionally, incomplete architectural recovery was visible at certain genomic loci where increased loop strength was maintained throughout the recovery period (Fig. 5c, Extended Data Fig. 6c). Importantly, this occurred without any detectable, permanent changes in the histone modification landscape. Finally, we found that globally, while differential loops that lost strength in TSA were fully restored, loops that became stronger upon TSA treatment remained enhanced (Fig. 5d).

**Figure 5.**
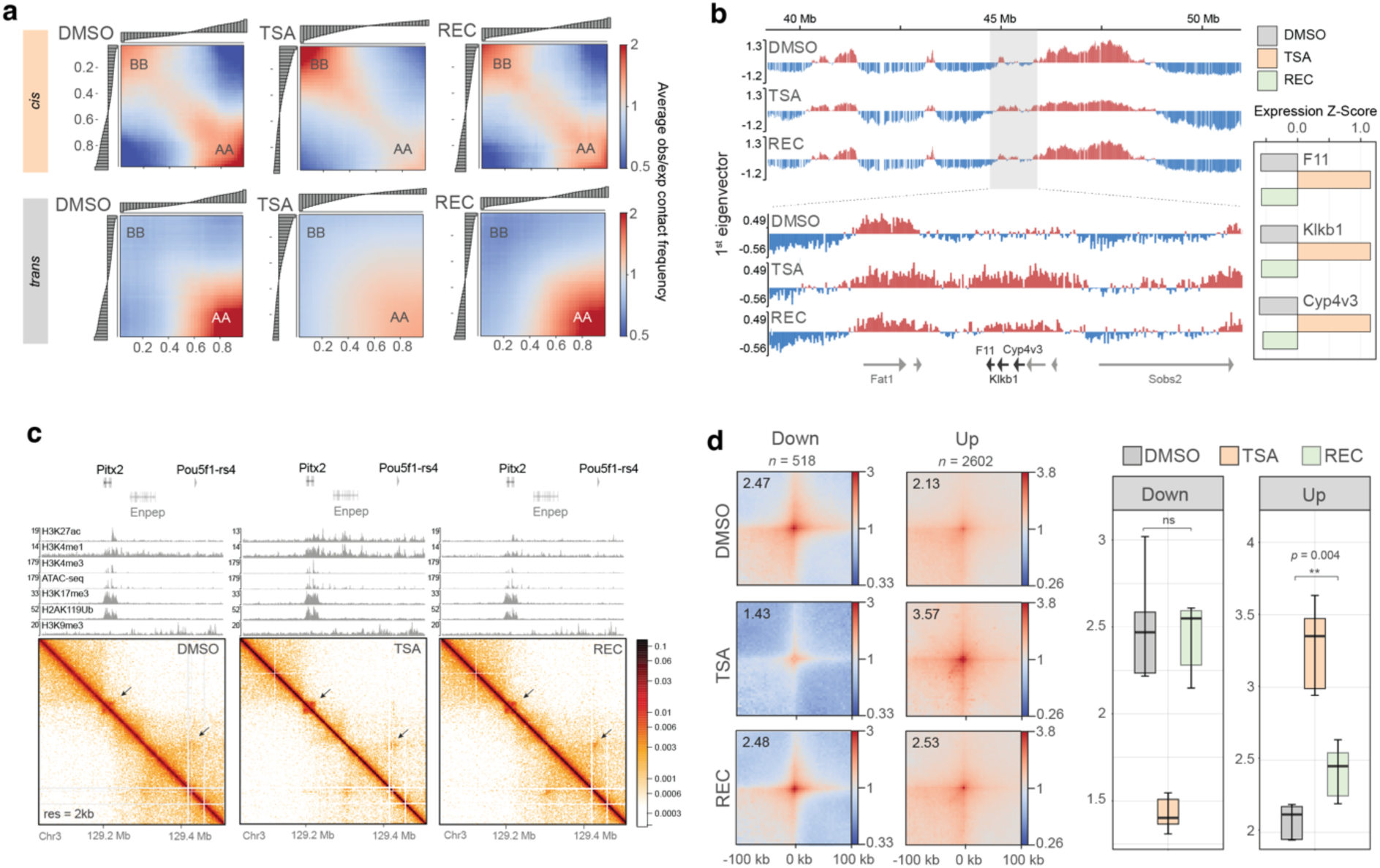
Genome architecture retains partial memory of the past conformation. **a**, Saddle plots of compartment interactions in *cis* (top panels) and in *trans* (bottom panels). **b**, High-resolution eigenvector tracks of the Micro-C data showing small-scale compartment switch around the F11-Klkb1-Cyp4v3 loci that persists in recovery. Gene expression Z-Score is shown in the right. **c**, Micro-C maps at the Pitx2 locus showing incomplete architectural recovery. ChIP-seq tracks of the corresponding condition are shown above. **d**, Pile-ups of Micro-C signal around differential loops in DMSO, TSA and recovery (resolution = 4 kb). Quantification of piled-up loop signal is shown on the right (paired two-tailed t-test; ns > 0.05, **p < 0.01). Data shown are the median, with hinges corresponding to IQR and whiskers extending to the lowest and highest values within 1.5× IQR.

In sum, we found that genome architecture carries a memory of its TSA-induced conformation that is visible at the level of *cis* contact frequencies, compartment interactions and at submegabase genome organisation.

### Sustained gene expression deregulation is associated with strong regulatory 3D contacts

We next reasoned that if the persisting minor architectural and transcriptional changes signified cellular memory, then repeated exposure to TSA should have more severe consequences. To test this hypothesis, we subjected mESCs to a second cycle of TSA treatment and recovery (Fig. 6a) which, similarly to the first, did not compromise cell viability or cell cycle progression (Extended Data Fig. 1a-d). Crucially, while the effects of the second TSA treatment were comparable to the first (Extended Data Fig. 7a, b), recovery from the second treatment was less complete (Fig. 6b). Hundreds of genes remained strongly deregulated (*n* = 767) and showed association with developmental processes, suggesting that the second pulse of perturbation had a greater impact on cellular identity (Fig. 6c). Next, we stratified differentially expressed genes based on their ability to recover from either TSA treatments, and we plotted their expression over the double treatment course. This revealed that, while most deregulated genes oscillated between their native and ectopic expression states, a subset of genes showed progressively aggravating gene expression deregulation (Fig. 6d), strongly indicative of cellular memory.

**Figure 6.**
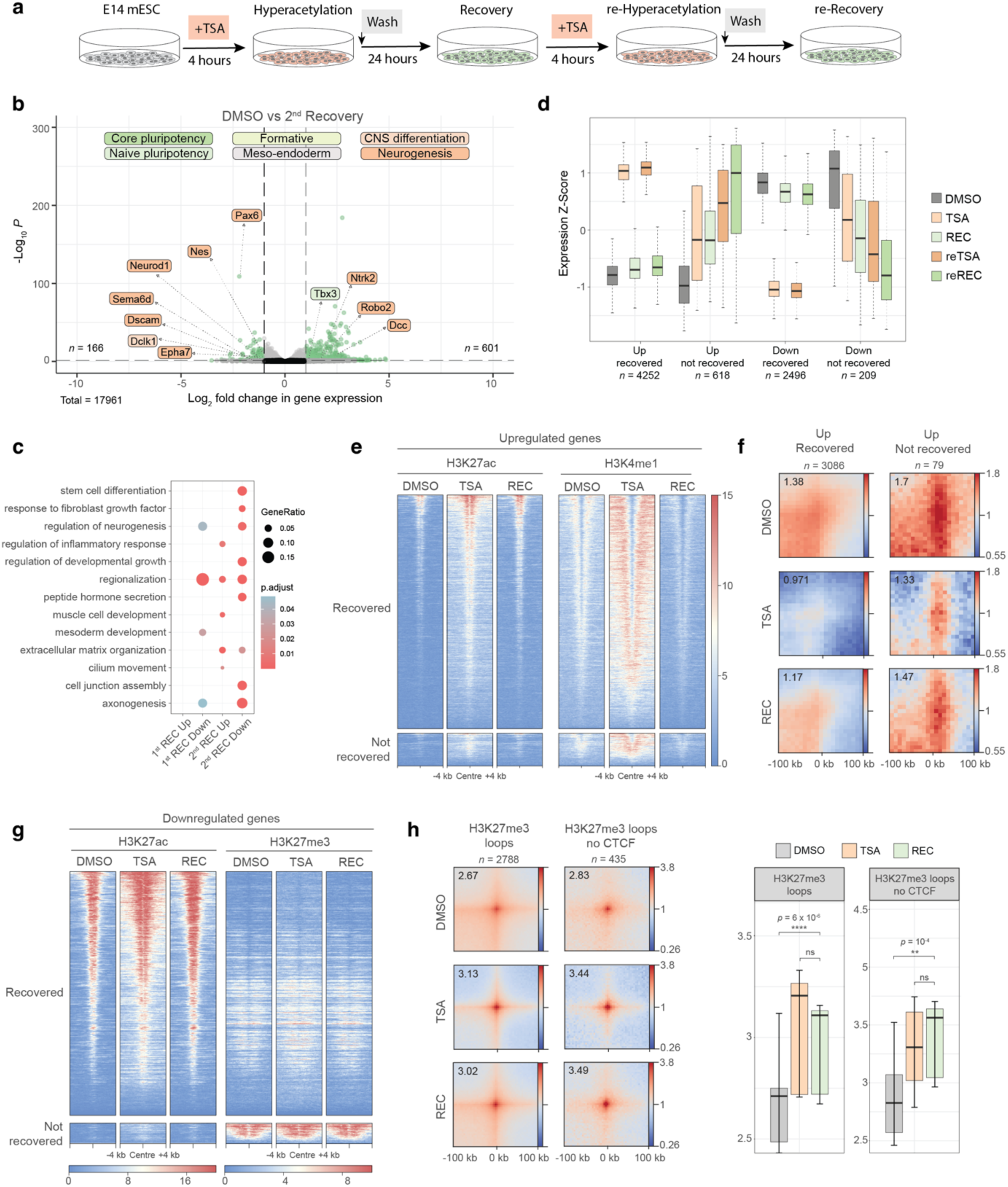
Sustained gene expression deregulation is associated with strong regulatory 3D contacts. **a**, Following the recovery period, cells were exposed to a second TSA pulse, wash and recovery cycle. **b**, Volcano plot showing differential gene expression (significance cutoffs: adjusted *p*-value > 0.05, absolute log_2_ fold change > 1) upon a second TSA (reTSA) treatment. Labels correspond to core and naïve pluripotency, formative, meso-endodermal, CNS differentiation and neurogenesis marker genes. **c**, Development-related Gene Ontology term enrichment among genes that remain misregulated following the first and second recoveries from TSA treatment. **d**, Gene expression Z-Scores of recovered and not recovered genes through the TSA-recovery treatment course. Data shown are the median, with hinges corresponding to IQR and whiskers extending to the lowest and highest values within 1.5× IQR. **e**, H3K27ac and H3K4me1 ChIP-seq signal in DMSO, TSA, and Recovery at upregulated TSS that recover (top) and do not recover (bottom). **f**, Pile-up of enhancer-promoter contacts in DMSO, TSA and Recovery around upregulated TSS that recover (top) and do not recover (bottom). **g**, H3K27ac and H3K27me3 ChIP-seq signal in DMSO, TSA, and Recovery at downregulated TSS that recover (top) and do not recover (bottom). **h**, Aggregate plots Micro-C signal in DMSO, TSA and Recovery at CTCF (left) and non-CTCF (right) loops with H3K27me3 ChIP-seq signal at loops anchors (resolution = 4 kb). Quantification of piled-up loop strength is shown on the right (paired two-tailed t-test; ns > 0.05, **p < 0.01. ****p < 0.0001). Data shown are the median, with hinges corresponding to IQR and whiskers extending to the lowest and highest values within 1.5× IQR.

Finally, we aimed to decipher what distinguishes genes that recover from genes that retain memory of their TSA-induced expression state. First, we analysed the histone modification landscape around upregulated TSSs. This analysis showed no particular difference between recovered genes and genes that did not recover: both groups were characterised by a gain of activating histone marks which were efficiently restored during the first recovery (Fig. 6e). Instead, a marked difference between the two gene groups was the presence of diffuse but strong pre-existing E-P contacts at non-recovery genes compared to recovery genes (Fig. 6f, Extended Data Fig. 7c). Next, similar TSS analyses among the downregulated genes revealed that genes that did not recover were Polycomb targets, as indicated by abundant H3K27me3 signal around promoters (Fig. 6g). Interestingly, downregulation happened without an apparent change in H3K27me3. Rather, we found that Polycomb loops gained substantial strength at non-recovery TSSs (Extended Data Fig. 7d-e), as well as genome wide, both at CTCF and CTCF-independent sites (Fig. 6h, Extended Data Fig. 7f). Critically, increased Polycomb-mediated looping persisted following the first recovery period, without change in H3K27me3 level at loop anchors (Extended Data Fig. 7h). Sustained loop strengthening was a feature specific to Polycomb rather than repressive loops in general, as we found the *de novo* H3K9me3 loops to recover efficiently (Extended Data Fig. 7g).

In conclusion, we found that repeated transient HDAC inhibition triggered cellular memory of gene expression at a subset of genes. We associate memory with strong architectural features surrounding deregulated TSSs: in case of upregulated genes prominent pre-formed E-P contacts, at downregulated TSSs bolstered repressive Polycomb loops perpetuate altered activity states. Importantly, these findings reveal a novel link between cellular memory and genome architecture.

## Discussion

The interplay between epigenetic layers and whether they function synergistically or antagonistically remains an area of active research. The findings presented in this study describe a crosstalk between activating and repressive epigenetic modifications, as well as genome folding, that profoundly modulate the mESC transcriptional programme. Namely, we uncover how acute perturbation of the histone acetylation landscape rapidly translates to changes in the histone methylation landscape and 3D chromatin organisation. Although the majority of epigenomic changes are reversible, we find that certain alterations in 3D genome folding persist and associate to a transcriptional memory effect at a subset of genomic loci.

Besides the general opening and activation of chromatin at promoters, we find widespread H3K4me1 deposition upon TSA treatment, indicating the deployment of new enhancers. These are likely to be major drivers of gene upregulation as previous studies have shown that the enhancer landscape – rather than promoter activity – is more significant for lineage determination^34,35^. Accordingly, enhancers are the most epigenetically dynamic regions of the genome^36,37^, explaining their susceptibility to the disruption of chromatin state balance. Interestingly, we find that H3K27 acetylation can trigger H3K4me1 deposition, which questions the commonly accepted sequence of events in enhancer activation where H3K4me1 is supposed to precede H3K27ac^38,39^. Once activated, the maintenance of enhancer activity is an active process^40–42^, explaining the efficient recovery of the enhancer landscape and transcriptional programme once the acetylation landscape is restored. Interestingly, we find that gene upregulation occurs without changes in E-P contacts, which agrees with recent studies that, in many instances, uncouple gene activation from a need to increase the frequency of physical contact^43–45^, or find that they are coupled only during terminal tissue differentiation but not in cell-state transitions^46^. Instead, we find pre-existing E-P contacts important for the memory effect. Indeed, it is thought that pre-formed E-P contacts may prime some genes for activation^20,25,47–50^, but additional triggers are required for transcription to take place. We speculate that excess H3K27ac activates enhancers, and those that are structurally in a high contact probability with their promoter targets can maintain active transcription even after the removal of ectopic acetylation.

Chromatin looping is commonly discussed in the context E-P contacts, or as the interplay between CTCF and cohesin that gives rise to loop extrusion-driven interactions. Our study highlights the importance and the potency of a third - often overlooked - type of focal contact corresponding to repressive chromatin loops. Somewhat counterintuitively, we find that excess chromatin activation and opening reinforces looping between loci marked by repressive chromatin signatures. One such class of loops corresponds to Polycomb (PcG) contacts that seem to be central to sustained gene expression downregulation. In neural progenitors PcG loci are known to exhibit transcriptional memory in *cis*, and this memory is linked to antagonism between Polycomb Repressive Complex 2 (PRC2) and activating signals^51^. Thus, one possible explanation is that excess genome-wide chromatin activation dilutes activating complexes away from PcG targets, tipping the balance towards gene downregulation. Crucially, as continued gene downregulation involves minimal – if any – change in the H3K27me3-H3K27ac balance at promoters, enhanced spatial sequestration of PcG loci appears to be central to the mechanisms of repression, constituting an architecture-based memory of a repressive state. Indeed, besides local chromatin compaction^52,53^, long range contacts are thought to be a mechanism by which PcG complexes confer silencing both in *Drosophila* and in mammals^48,54–58^, thus it is plausible that chromatin organisation would also be a mechanism of PcG-driven epigenetic memory^59^. We also detect prominent looping between H3K9me3 marked loci as a potential mechanism of gene downregulation which, contrarily to PcG loops, does not show memory. In the future it might be interesting to investigate if H3K9me3 contacts are mediated by homotypic chromatin interactions, or if looping involves the action of chromatin binding proteins such as HP1^60^, perhaps with specific associated partners that might gain interactions upon changes in chromatin acetylation state.

Other than fine-scale changes precisely linked to differences in gene expression, acute disruption of the acetylation landscape led to important changes in global genome folding. The increase in *trans* contacts points to the possibility that histone acetylation might be an important determinant of intrachromosomal interactions as well as chromosome territories. Indeed, it has been shown that long, highly transcribed genes or gene dense regions extend from chromosome territories^21,61–63^, although this has been attributed to binding of ribonucleoproteins to nascent transcripts rather than to the acetylation state *per se*. Using biophysical modelling we found that, by increasing the stiffness of the chromatin fibre^21^, we can indeed accurately recapitulate the *trans* contact ratio observed in TSA. While the determinants of *trans* contacts remain largely elusive, it is widely accepted that homotypic interactions between domains of the same epigenetic state are the major driving force of chromosome compartmentalisation^64^. Thus, it was reasonable to anticipate that histone hyperacetylation would have an impact on A and B compartments. However, to our surprise global chromatin activation weakened rather than strengthened A-A compartment interactions to an extent that is comparable to what occurs during ESC differentiation^20^. Interestingly, simulations that we carried out in an attempt to understand whether increased chromatin stiffness can give rise to excess *trans* contacts efficiently predicted the change in compartment interactions that we observed in the Micro-C data. This suggests that chromatin stiffness is an important biophysical determinant of not just interchromosomal contacts but also A/B compartmentalisation.

Finally, we found that these perturbed compartment interactions can partially persist beyond the recovery period, signifying that 3D structures carry a partial memory of their past state. This might be explained with hysteresis, the dependence of a system’s behaviour on its history. Hysteresis is an emerging principle in 3D genome organisation which was found to be critical to model certain characteristics of genome folding, ranging from *cis* contacts in gene expression control^65^, to organisation of the interphase nucleus^66^. Such a principle has been demonstrated to exist by experimental approaches, whereby interphase chromosome conformation was found to depend on the condensin complex that carried out mitotic chromosome condensation in the previous cell cycle^67^. Our study is in line with these observations and provides empirical evidence of an architectural memory both at global scale and at gene level. Biophysical modelling has shown that 3D genome folding might be a critical element to stabilise epigenetic memory in interphase cells, providing further support to association that we found between 3D genome folding and cellular memory^68–70^. In the future it will be critical to understand how mitotic events in the nucleus might be used by cells to modulate or maintain their identity, as well as what molecular factors contribute to this process.

## Data Availability

All raw data were submitted to the National Library of Medicine’s (NCBI) Sequence Read Archive (SRA) and the processed files were submitted to the Gene Expression Omnibus (GEO). All data can be retrieved under the GEO SuperSeries GSE281151. Micro-C pairing statistics are detailed in Supplementary Table 1. Differential gene expression and differential chromatin looping results can be found in Supplementary Table 2. ChIP-seq, RNA-seq and loop anchor Gene Ontology enrichment can be found in Supplementary Table 3.

## Code Availability

Scripts used for biophysical modelling in this article are available at https://github.com/cavallifly/Paldi_et_al_2024.

## Acknowledgements

We thank Ivana Jerkovic and Frederic Bantignies for comments on the manuscript. We are grateful to Dovetail Genomics (Cantata Bio) for their contribution to Micro-C reagent costs. We are grateful to the Genotoul bioinformatics platform Toulouse Occitanie (BioinfoGenotoul, https://doi.org/10.15454/1.5572369328961167E12) and the Pôle Scientifique de Modélisation Numérique (PSMN) of the ENS de Lyon for computational resources. We thank the Montpellier Resources Imagerie (MRI) for their help and support with microscopy. FP was supported by HFSP Long-term Fellowship LT000111/2021-L and EMBO Long-term Fellowship ATLF 716-2020. MS was supported by a Marie Skłodowska-Curie Innovative Training Network (grant no. 813327 ‘ChromDesign’) and Fondation de Recherche Medicale. HR was supported by La Ligue Contre le Cancer. This work in the Cavalli lab was funded through grants from the European Research Council (Advanced Grant 3DEpi), from the “Agence Nationale pour la Recherche” (EpiGenMed ANR-10-LABX-12-01; PLASMADIFF3D, grant N. ANR-18-CE15-0010, LIVCHROM, grant N. ANR-21-CE45-0011), by the European E-RARE NEURO DISEASES grant “IMPACT”, by the Fondation ARC (EpiMM3D), by the Fondation pour la Recherche Médicale (EQU202303016280), by the MSD Avenir Foundation (Project GENE-IGH), and by the French National Cancer Institute (INCa, PIT-MM grant N. INCA-PLBIO18-362).

## Contributions

GC and FP conceived the study. GC supervised the project and FP designed and carried out wet-lab experiments and analysed all data. MS optimised gastruloid culture, and helped to carry out FACS, ChIP-seq and ATAC-seq experiments as well as Micro-C data analysis. MDS and DJ designed biophysical modelling. MDS carried out polymer simulations and analysis. HR performed YY1 ChIP-seq. FP wrote the manuscript with input from all authors.

## Competing Interests

The authors declare no competing interests.

## Material and Methods

## EXPERIMENTAL METHOD DETAILS

### Embryonic stem cell culture

E14GT2a p14 cells were purchased from MMRRC, UC Davis, CTCF-GFP-AID cells were gift from R. Saldana-Meyer. ESCs were cultured on plastic plates coated with 0.1% gelatin (Sigma-Aldrich #G1890-100G) in serum-LIF medium (GMEM (Gibco #2171002), with 15% FBS (ThermoFisher #26140079 USA origin), 1x Glutamax (ThermoFisher #35050038), 1x MEM Non-Essential Amino Acids (ThermoFisher #11140035), 50 U Penicillin-Streptomycin (Gibco #15140122), 0.1 mM Sodium Pyruvate (Gibco #11360070), 0.1 mM 2-Mercaptoethanol (Gibco #31350010), and 1,000 U/ml LIF (Sigma-Aldrich #ESG1107)). Cells were passaged every 2-3 days using TrypLE Express Enzyme (Gibco #12604013). Cell lines were regularly tested for mycoplasma infection. Cell viability was assessed by staining with Trypan Blue (Gibco # 15250061) and cells were counted on a Countess 3 automated cell counter (Invitrogen). HDAC inhibition was performed by treating cells with 100 ng/ml Trichostatin A (Sigma-Aldrich #647925) for 4 hours. Control cells were treated with 0.01% DMSO for the same duration. For recovery, cells were washed once with PBS and were incubated with fresh mESC media for 10 minutes. This PBS wash/media change was repeated 2 more times before incubating cell for a total of 24 hours.

### Gastruloid culture

Gastruloids for RNA-seq and immunostaining experiments were generated as described in^71^. Briefly, CTCF-GFP-AID cells were harvested, centrifuged and washed twice with PBS. Following, cells were resuspended in N2B27 medium and counted. 300 cells were seeded in each well of a round-bottomed, low-attachment 96-well plate (Greiner #650970) in N2B27 medium. After 48 hours, a 24-hour pulse of 3 µM Chiron was administered and media was changed every day. HDAC inhibition was carried out by treating gastruloids with 20 ng/ml ml Trichostatin A (Sigma-Aldrich #647925) for 4 hours immediately prior to the Chiron pulse (44-48 hours). TSA was removed from the medium by changing N2B27 media three times with 10 minutes of incubation in between. Control cells were washed similarly.

### Western blotting

For western blotting ∼10^7^ mESCs were dissociated, washed once in PBS, resuspended in 200 *μ*l of Cell Lysis Buffer (85mM KCl; 0,5% NP40; 5mM HEPES pH. 8; 1X EDTA-free Protease inhibitor (Roche); 5mM sodium butyrate) and incubated on ice for 15 minutes. Afterwards, cell nuclei were pelleted at 2000g for 5 minutes at 4°C. The supernatant (cytoplasmic fraction) was separated, and nuclei were resuspended in 100 *μ*l RIPA buffer (50 mM Tris pH 7.5; 150 mM NaCl; 1% NP40; 0,5 % NaDoc; 0,1% SDS; 1X EDTA-free Protease inhibitor (Roche #04693132001); 5mM sodium butyrate). Following a 10-minute incubation on ice, chromatin was digested for 15 min at 37°C with 0.0125 U/μl MNase and 1mM of CaCl_2_. Extracts were cleared by 30 minutes of centrifugation at 14000 rpm at 4°C. Protein yield was quantified using Pierce BCA protein assay kit (ThermoFisher #A65453). Samples were mixed with 4x NuPage LDS sample buffer (ThemoFisher #NP0007) and boiled for 10 minutes at 95°C. 2 µg denatured protein extract was loaded per lane on a NuPAGE™ 4-12 %, Bis-Tris gel (ThermoFisher #NP0321BOX). Transfer onto nitrocellulose membranes was performed using the Trans-Blot Turbo Transfer System (Biorad). Membranes were stained with Ponceau S for 5 minutes, then blocked for at least 30 min with 3% BSA in PBS + 0.1% Triton-X100 prior to incubation with primary antibody overnight at 4°C with the following dilutions: α-H3K27ac 1:7500 (Active Motif #39133); α-pan-acetyl lysine 1:1000 (ThermoFisher, #66289-1-IG); α-lamin B1 1:10000 (abcam #ab16048); α-Vinculin 1:1000 (Santa Cruz Biotechnology #sc-73614). Membranes were washed three times >5 minutes in PBS +0.1% Tween-20 and were incubated with secondary antibodies (α-Rabbit IgG-Peroxidase antibody (Sigma-Aldrich #A0545) or α-Mouse IgG–Peroxidase antibody (Sigma-Aldrich #A9044)) at 1:16000 dilution for 1 hour at room temperature. After three >5-minute washes with PBS-0.1% Tween-20 at room temperature, membranes were developed using the SuperSignal West Dura Extended Duration Substrate solution (ThermoFisher #34075) for 1 minute and imaged with a Bio-Rad ChemiDoc imager.

### Flow Cytometry

1-3×10^6^ mESCs were dissociated with TrypLE, pelleted, and resuspended in PBS. For cell cycle analysis, dissociated mESCs were washed once in PBS and pelleted and fixed in cold 70% ethanol for 30 min at 4°C. Cells were stained with the Propidium Iodide Flow Cytometry Kit (Abcam #ab139418) according to manufacturer’s instruction. Flow cytometry was performed on a Miltenyi MACSQuant instrument, and analysis was performed using the FlowJo software.

### Gastruloid immunostaining

Gastruloid immunostaining protocol was adopted from^72^. Plastic material was pre-coated with the blocking solution (PBS + 10% FBS + 0.2% Triton-X100). Using a cut P1000 tip, gastruloids were collected into 15 mL centrifuge tubes. Following a PBS wash, gastruloids were transferred to with 2mL 4% PFA in 6-well plates and were fixed overnight at 4°C. For washes, gastruloids were transferred serially across three PBS-filled wells and were incubated for 10 minutes in the last one. Gastruloids were blocked in PBS+FT (PBS + 10% FBS + 0.2% Triton-X100) for 1 hour at room temperature, then incubated with primary antibodies (α-Brachyury 1:500 (Santa Cruz Biotechnologies #sc-166962), α-Sox2 1:500 (eBioscience #15208187), α-H3K27ac 1:200 (Active Motif #39133)) in PBS+FT and 1 ug/ml DAPI overnight at 4°C with orbital shaking. Gastruloids were washed by sequentially transferring them in 3 wells filled with PBS+FT and incubating them for 20 minutes in the last one. Staining with secondary antibody (α-Rabbit Alexa Fluor Plus 488 1:400 (ThermoFisher #A32731); α-Rabbit Alexa Fluor Plus 555 1:400 (ThermoFisher #A32794); α-Rat Alexa Fluor Plus 647 1:400 (ThermoFisher #A48265)), and 1 ug/ml DAPI, as well as washes were carried out similarly to primary antibody. Gastruloids were mounted in ∼30uL of Fluoromount-G (ThermoFisher #00-4958-02) and were kept at 4 °C before imaging.

### Image acquisition and quantification

Confocal imaging was performed using a Confocal Zeiss LSM980 Airyscan II equipped with a 20x objective. Diodes laser 405, 488, 561 and 639 nm were used for fluorophore excitations, leading to blue, green, red and far-red channels. For each gastruloid 3 z-stacks were taken and using the Fiji software, maximum intensities were projected to manually define areas of H3K27ac, Sox2 and Brachyury expression as well as DAPI staining.

### RNA isolation for RNA-seq

RNA was isolated using the RNeasy mini kit (Qiagen 74104). Cells were detached with TrypLE, lysed in RLT buffer with β-mercaptoethanol and lysates were processed according to manufacturer’s instruction. For mESCs columns and buffers supplied with the RNeasy kit were used, while for gastruloids the Zymo RNA Clean & Concentrator-5 (Zymo Research #R1015) reagents were used. On-column DNAse-I digestion (Qiagen #79254) was performed as recommended. RNA samples were sent to BGI Tech Solutions (Hongkong) for strand-specific transcriptome sequencing. Samples were sequenced at a depth of 50 million 150 bp paired end reads. All experiments were performed in triplicates.

### Micro-C library preparation and sequencing

Micro-C libraries were generated with the Dovetail™ Micro-C Kit protocol v1.0 with minor modifications. Briefly, 10^6^ mESC were washed with PBS and were frozen at −80°C for at least 1 hour. Cell pellets were thawed and crosslinked first with 3 mM DSG (ThermoFisher #A35392) in PBS for 10 minutes at RT with rotation, then formaldehyde was added at 1% final concentration for further 10 minutes. The pellets were washed twice with PBS and digested with MNase according to kit instructions. MNase digestion was routinely verified by decrosslinking a small amount of chromatin and assessing fragment distribution on a Bioanalyzer 2100 instrument (Agilent). If the digestion profile showed 60% - 70% mononucleosomal DNA fraction, on-bead proximity ligation was performed, followed by reversed cross-linking and DNA purification. End repair and adapter ligation were performed using the NEBNext Ultra II DNA Library Prep Kit for Illumina (NEB #E7645). Following, DNA was purified using SPRI beads (Beckman #B23318) as described in the Micro-C user manual. Finally, biotin pulldown and library amplification were performed according to Dovetail Micro-C Kit User Guide and using Dovetail Micro-C Kit reagents, only replacing the Dovetail Primers (Universal and Index) with NEBNext primers. Libraries were pooled and sent to BGI Tech Solutions (Hongkong) for 100-bp paired-end sequencing to obtain roughly 2-3 billion reads for each replicate in the present study.

### Chromatin Immunoprecipitation followed by sequencing (ChIP-seq)

Chromatin immunoprecipitation was performed as described previously^73^. Cells were collected with TrypLE (ThermFisher #12604013) and fixed with 1% methanol-free formaldehyde in mESC media for 10 minutes with rotation at RT. Glycine (2.5M Glycine in PBS) was used to stop the fixation for 10 minutes with rotation at RT. Fixed cells were centrifuged at 500 g for 5 minutes at 4°C, washed twice in 1x ice-cold PBS and snap frozen in liquid nitrogen until further use. Following thawing, cells were spiked-in with 8% HEK-293 cells and chromatin extraction was performed as in^74^. 15 μg of chromatin was used for each replicate of histone ChIP, and 50 µg for CTCF and YY1 ChIP, with 6–8 μg of antibody. Since the above protocol was not suitable for YY1, we followed the protocol described by^75^. Briefly, fixed cells resuspended in SDS buffer, followed by sonication and preparation for immunoprecipitation in a homemade buffer. Next, the mixture was incubated overnight at 4°C with Protein G beads (Invitrogen #10004D), washed with both low and high-salt buffers, reverse-crosslinked in an elution buffer, and purified using a QIAQuick PCR purification kit (Qiagen # 28104). Antibodies used in this study were: H3K4me1 Active Motif #39297), H3K4me3 (Milipore #04-745), H3K27ac (Active Motif #39133), H3K9me3 (abcam ab8898), H3K27me3 (Active Motif #39155), H2AK119Ub (Cell Signalling #8240S), CTCF (Active Motif #61311), and YY1 (abcam 109237). Sequencing libraries were constructed using NEBNext Ultra II DNA Library Prep Kit for Illumina (NEB #E7645), pooled and sent to BGI Tech Solutions (Hongkong) for 100-bp paired-end sequencing to obtain roughly 30-50 million reads for each replicate. All experiments were performed in duplicates.

### Assay for transposase accessible chromatin (ATAC-seq)

For each replicate, 9×10^4^ mESCs were collected with TrypLE (ThermFisher # 12604013) and were mixed with 10^4^ HEK-293 cells. Samples were processed using the Active Motif ATAC-seq kit (Active Motif #53150) following manufacturer’s instructions without modifications. ATAC-seq libraries were pooled and sent to BGI Tech Solutions (Hongkong) for 100-bp paired-end sequencing to obtain roughly 30-50 million reads for each replicate. All experiments were performed in duplicates.

## QUANTIFICATION AND STATISTICAL ANALYSES

### RNA-seq analysis

RNA-seq samples were mapped using the *align* function of the Subread package (2.0.6). Subread command *featureCounts* (with options “*-p --countReadPairs -s 2 -t exon*”) and the feature file UCSC RefSeq GTF file for mm10 were used to generate count tables that were then used as an input for the DEseq2^76^ to perform the differential analysis (Supplementary Table 2). Gene Ontology analysis was performed significantly up- and down-regulated genes (p-adjust ≤ 0.05, |log2FC| ≤ 1) using the *enrichGO* function from the clusterProfiler package^77^ (Supplementary Table 3). Volcano plots and scatter plots were produced in R using the EnhancedVolcano and ggplot2 libraries respectively.

### ChIP-seq and ATAC-seq analysis

ChIP-seq and ATAC-seq samples were mapped using bowtie2 v.2.4.4^78^ with command “*bowtie2 -p 12 --no-mixed --no-discordan*t” against the mm10 and hg19 genomes. Then, samtools v.1.9^79^ was used to filter out low-quality reads (command “*samtools view -b -q 30*“) and Sambamba v1.0^80^ was used to sort (command *“sambamba sort”*), deduplicate and index bam files (*“sambamba markdup --remove-duplicates”*) with default parameters. Following, samtools was used to count both human a mouse reads (command “*samtools view -c”*) to calculate down-sampling factor (dF) for spike-in normalisation as described in^81^. Next, bam files were downscaled accordingly using samtools (command “*samtools view -b -s dF”*) and bigwig files were produced using the deepTools package^82^ with command “*bamCoverage -- normalizeUsing none --ignoreDuplicates -e 0 -bs 10*”. Finally, ChIP-seq tracks in were visualized using the IGV v.2.16.1 software^83^ or HiGlass^84^. ATAC-seq and ChIP-seq peaks were called on each replicate using MACS3 with a q value cutoff 0.05, and for histone marks with the additional parameters *--broad --broad-cutoff 0.1*s^85^. Finally, peaks detected from both replicates were filtered and all downstream analyses were carried out using this consensus peak set. For differential peak calling the diffBind^86^ R package was used with normalisation “*normalize=DBA_NORM_LIB, spikein = TRUE*”, analysis method “*method=DBA_DESEQ2*”, and FDR < 0.05 cutoff. Heatmaps and metaplots were produced using the *computeMatrix* function of the deepTools package, and plotted using the *plotHeatmap* and *plotProfile* functions. ChIP-seq bloxplots were also created by deepTools using the the *multiBigwigSummary* function and were plotted by ggplot2 in R. Chromosome wide H3K27ac read density plots were generated using a custom R script published in^81^. ChIP-seq peak distribution and annotation were carried out with ChIPseeker’s^87^ *plotPeakProf* and *annotatePeak* functions, respectively. Gene ontology analysis of annotated ChIP-seq peaks was performed using the *enrichGO* function from the clusterProfiler package^77^ (Supplementary Table 3). For cumulative histograms, enhancer distance from TSSs was calculated using bedtools closest function and was plotted by ggplot2 in R. Expression-matched control gene set was derived using code from the AdelmanLab github repository (https://github.com/AdelmanLab/Expression-Matching). Myc ChIP-seq dataset was published in^88^ and was downloaded from the GEO repository GSE90895.

### Micro-C data analysis

#### Generation of contact matrices and standard analyses

Micro-C data was mapped using the HiC-Pro v3.1.0 pipeline^89^. Fastq reads were trimmed to 50 bp using TrimGalore (-- hardtrim5 50) (https://github.com/FelixKrueger/TrimGalore) and aligned to the mm10 (UCSC Mouse GRCm38/mm10) reference genome using bowtie2^78^ v2.4.4 *(“--very-sensitive --L 30 -- score-min L, -0.6, −0.2 --end-to-end --reorder*”), removing singleton, multi-hit and duplicated reads. Minimum *cis* distance was set at 200 bp and non-informative pairs were removed. The total numbers of valid read-pairs per sample are reported in Supplementary Table 1. Contact matrices in the .cool file format were generated using cooler^90^ v.0.10.2 at 100 bp resolution (command “*cooler cload pairs -c1 2 -p1 3 -c2 5 -p2 6 ./scripts/chrom_sizes.txt:100*”). Similarities between replicates (5 replicates for DMSO and TSA; 2 replicates for 24-hour recovery) were measured using the <u>S</u>tratum-adjusted <u>C</u>orrelation <u>C</u>oefficient (SCC) applying *HiCRep* v1.12. (https://github.com/TaoYang-dev/hicrep)^91^ on chromosomes 2, 9, 13, and 19 using the *get.scc* function with parameters resol=20kb and (lbr,ubr,h)=((0, 100kb,1), (100kb, 500kb,1), (500kb, 2Mb,2), (2Mb, 10Mb,4)). H values were previously trained using the *htrain()* on two replicates (E and F) of the DMSO condition. Using 1.0-SCC, as a measure of the similarity (0 - similar and 1 dissimilar) between replicates and hierarchical clustering analysis using *hclust()* function in R with Ward.D2 method on the chromosome-averaged similarities, allowed us to distinguish and group together the replicates of the different conditions, motivating us to merge the valid-pairs of different replicates in a unique dataset for each condition. Multiresolution *.mcool* files were obtained and normalized via the Iterative <u>C</u>orrection and <u>E</u>igenvector decomposition algorithm (ICE) with default parameters (command “*cooler zoomify -r 100, 400, 600, 800, 1000, 2000, 4000, 8000, 10000, 20000, 50000, 100000, 1000000, 10000000 file.cool -o file.mcool --balance*”)^92^ and were uploaded onto a local HiGlass server for visualization^84^. For comparison of architectural features between different conditions contact maps were matched to contain approximately the same number of *cis* contacts (Supplementary Table 1). All genomic snapshots of Micro-C maps were generated using HiGlass v.1.11.7. Standard analyses (P-value curves, eigenvector analyses, saddle plots, insulation score analysis) were performed using the cooltools (version 0.5.4) package. **Loop analyses**. Loops were called using mustache v1.0^93^ with default parameters *(“-- pThreshold 0.1 -–sparsityThreshold 0.88 -–octaves 2*”) on ICE-balanced maps at 1 kb and 4 kb resolutions. Redundant loops between different resolutions were filtered in 20 kb windows, and in case of overlap loop coordinates were retained at the finer resolution. All aggregate plots were created with the coolpuppy v1.1.0 package^94^ and were normalised using expected maps generated by cooltools. For differential looping, contacts that overlapped with the corresponding loop anchor bin were summed for each loop and were summarized into a count table: genome-wide count tables were created for each replicate at each resolution (command *“cooler dump --join -t pixels”*) then filtered against loop using bedtools ‘pairtopair’ function^95^. The count tables from different conditions were used for differential analysis with DESeq2 (Supplementary Table 2). The thresholds p.adj < 0.05, |log2FoldChange| > 0.5 and baseMean >= 10 were used to filter for significant changes in looping between conditions. Volcano plots were produced in R using the EnhancedVolcano library. Loop subclasses were defined based on the presence of ChIP-seq peaks at loop anchors (repressive: overlapping with H3K9me3, H3K27me3 or H2AK119Ub peak; active: overlapping with H3K4me1, H3K4me3 or H3K27ac peak; *de novo* H3K9me3 loops: loops only present in TSA overlapping with H3K9me3 peak; CTCF: loop anchors within +/- 1kb of CTCF peaks; non-CTCF: no CTCF peak within +/- 2.5 kb of loop anchor) or presence of TSSs within 2 kb of either loop anchor. Enhancer-promoter contacts for recovery vs non-recovery genes were taken from^20^. Loop quantification boxplots represent the observed/expected value of the central 5×5 pixels of aggregate plots that was extracted from coolpuppy matrices using an in-house Python script. Loop anchors were annotated using the *annotatePeak* function of the ChIPseeker R package, and annotated anchors within <10 kb from TSSs were used for GO enrichment with the *enrichGO* function of clusterProfiler library (Supplementary Table 3).

### Biophysical modelling

Two polymeric systems were prepared in 30 and 5 replicates respectively with 1 and 20 chains of 20 mega-basepairs (Mbp) each. Each chain-bead was of unitary mass (m = 1.0), hosting *v* = 5 kilo-base pairs (kbp) of DNA sequence and has a diameter of σ. This representation was obtained with the Kremer-Grest bead-spring model^96^ with the same parameter as in^97^.

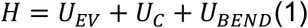

The first term was a truncated and shifted Lennard-Jones potential that controls the *cis*- and *trans*-chromosome excluded volume interactions:

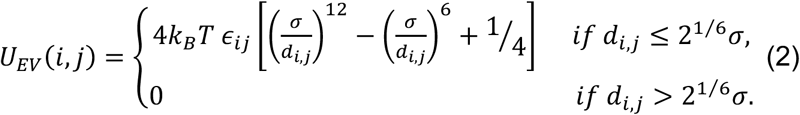

where k_B_ is the Boltzmann constant, T the temperature, *ϵ_ij_* is equal to 10 if |*i* − *j*| = 1, and 1 otherwise, σ was the thickness of the chain and d_i,j_ is the modulus of 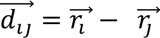, that is the distance vector between the monomers *i* and *j* at positions 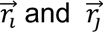, respectively.

The second term was a FENE potential that maintains chain connectivity between consecutive beads on the same polymer chain:

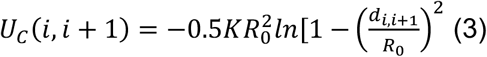

where K=0.33 k_B_T/nm^2^ and R_0_=1.5σ. The combined action of the connectivity and excluded volume interaction between consecutive beads was such that the average bond length was close to σ and never exceeded 1.1σ.

The third term is a (Kratky-Porod) bending potential:

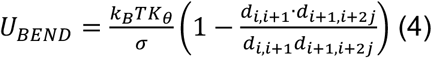

where *K*_θ_ is the chain persistence length.

The dynamics of the polymer model was simulated using the LAMMPS simulation package (version 29 Oct 2020) integrating the (underdamped) Langevin equation of motion^98^:

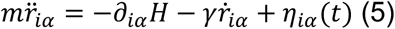

where m is the mass of the bead that was set equal to the LAMMPS default value, H is the Hamiltonian of the system in Eq. (1), the index *i* runs over all the particles in the system, and *α* = (*x*, *y*, *z*) indicates the Cartesian components, and γ = 0.5 τ_LJ_^−1^ is the friction coefficient with τ_LJ_ = σ(m/ε)^1/2^ is the Lennard-Jones time. The stochastic term *η*_*iα*_ satisfies the fluctuation-dissipation conditions. The integration time step used in the numerical integration was equal to Δt = α τ_LJ_, where the factor α was adapted, as specified below, to the different stages of the preparation and production runs.

#### Preparation of the initial conformations

Each chain is initially organized in a rod-like folding featuring rosettes along the main axis and placed in random positions inside a confining sphere of radius R* so to set up the volume density of the system to 3% (R*= 25.5σ for the 1-chain system and R*=69.3σ for the 20-chain system), avoiding clashes with other chains. The confining sphere is completely contained in a cubic simulation box with fixed boundary conditions. After an energy minimization (*LAMMPS command*: minimize 1.0e-4 1.0e-6 100000 100000), each of the polymeric system is compressed to reach the DNA density of 10%. These conditions were achieved by minimization (*LAMMPS command*: minimize 1.0e-4 1.0e-6 100000 100000) followed by molecular dynamics simulations of 600 τ_LJ_ (100,000 Δt with Δt = 0.006 τ_LJ_) during which the radius of confining spheres is reduced from the minimum radius to include all the particles of the chains at time 0 to the target radius R (R= 17.1σ for the 1-chain system and R=46.4σ for the 20-chain system). At the target volume density of 10%, the polymer chains have parameters σ∼54.2nm and *K_θ_* ∼92.3nm. These estimates were done by considering a fine-scale chromatin model with *v*_*FS*_ = 100 *bp*, σ_FS_ = 20 *nm* and *K_θFS_* = 50 *nm* and the coarse-grain procedure in^99^. Finally, each polymeric system is relaxed with molecular dynamics run of 30,600 τ_LJ_ (5,100,000 Δt with Δt = 0.006 τ_LJ_). By comparing the average monomer <u>M</u>ean-<u>S</u>quared <u>D</u>isplacement (MSD) in these relaxation runs and the MSD of non-transcribed genes measured^100^ by live-cell imaging, we obtained an approximated estimate of the simulated time (in τ_LJ_) corresponding to 1s ∼ 9 τ_LJ_. These conformations are next used as the initial conformations for the downstream simulations.

#### A/B compartmentalisation

To model the A/B compartmentalisation in the DMSO condition short-range interactions were used to test the attractions between the model regions which correspond to A/B compartments. As shown in Fig. 2d, the first and the last 1 Mb were assigned to telomeric region and 6 blocks of A/B domains each of 1.5 Mb were defined in the central part. These interactions have been modelled using attractive Lennard-Jones potentials (see **Equation 2**) with cutoff=2.5σ, that allowed to include the attractive part of the Lennard-Jones potential. To allow efficient parameter sampling, we performed 4h simulations (21,600,000 Δt with Δt = 0.006 τ_LJ_) for just one chain by varying the strengths of compartments’ interactions (E_AA_ and E_BB_) were varied in the range 0.00-0.40 k_B_T in the 1-chain system with 4h trajectories (21,600,000 Δt with Δt = 0.006 τ_LJ_). To infer these energies, A/B compartment-strengths at 10kb from the micro-C maps were matched against the correspondent quantities computed on the model chains^66^. The compartment strength (CS) profile is obtained by partitioning the A-(B-) domains in 150 bins and by averaging within each of them the CS per 10kb-bin of the Micro-C or models’ contact-maps. The distance-cutoff for detecting contacts in the models was set to 150 nm∼3beads. The Euclidean distance between A and B profiles was used to define compartment-specific ranks r_A_ and r_B_ ranging from 1 to max(r) where 1 is the best match with the experiments. A unique rank r was defined from the average of r_A_ and r_B_. Finally, the similarity score was defined as (r-max(r))/(min(r)-max(r)) and it is equal to 1 for the condition that best describes the experimental CS-profile and to 0 for the least accurate one. This procedure resulted in the optimal values E_AA_=0.080 and E_BB_=0.00. Next, the DMSO and TSA condition were modelled by the system made of 20 chains simulated for with 4h-trajectories (21,600,000 Δt with Δt = 0.006 τ_LJ_). The optimized A/B-compartment attractions (ε_AA_=0.080 and ε_BB_=0.00) were maintained and the bending rigidity of the A- and B-compartment domains were differentially increased. We explored several combinations varying *K_θ_* between 1 and 16 times and *K_θ_* between 0 and 5 times the nominal persistence length of *K_θ_* ∼92.3nm. The chromosome-averaged *trans*-contact ratios (TR) were computed by averaging the trans-contact ratio of 10kb-bins in each chromosome. The median was obtained on the distribution of these chromosome-averaged values. The absolute difference between the median values TR for all, only A and only B domains in the micro-C datasets and the models was used to define three ranks per each parameter set. A similarity score was defined from these three ranks applying a strategy analogous to one described above for CS profiles. Optimal values were (*K_θ_*, *K_θ_*) = (1,0)*K*_θ_ in DMSO and (*K_θ_*, *K_θ_*) = (14, 3)*K_θ_* in TSA. Model snapshots in Extended Data Fig. 2j were prepared using VMD^101^.

---

**Extended Data Figure 1.**
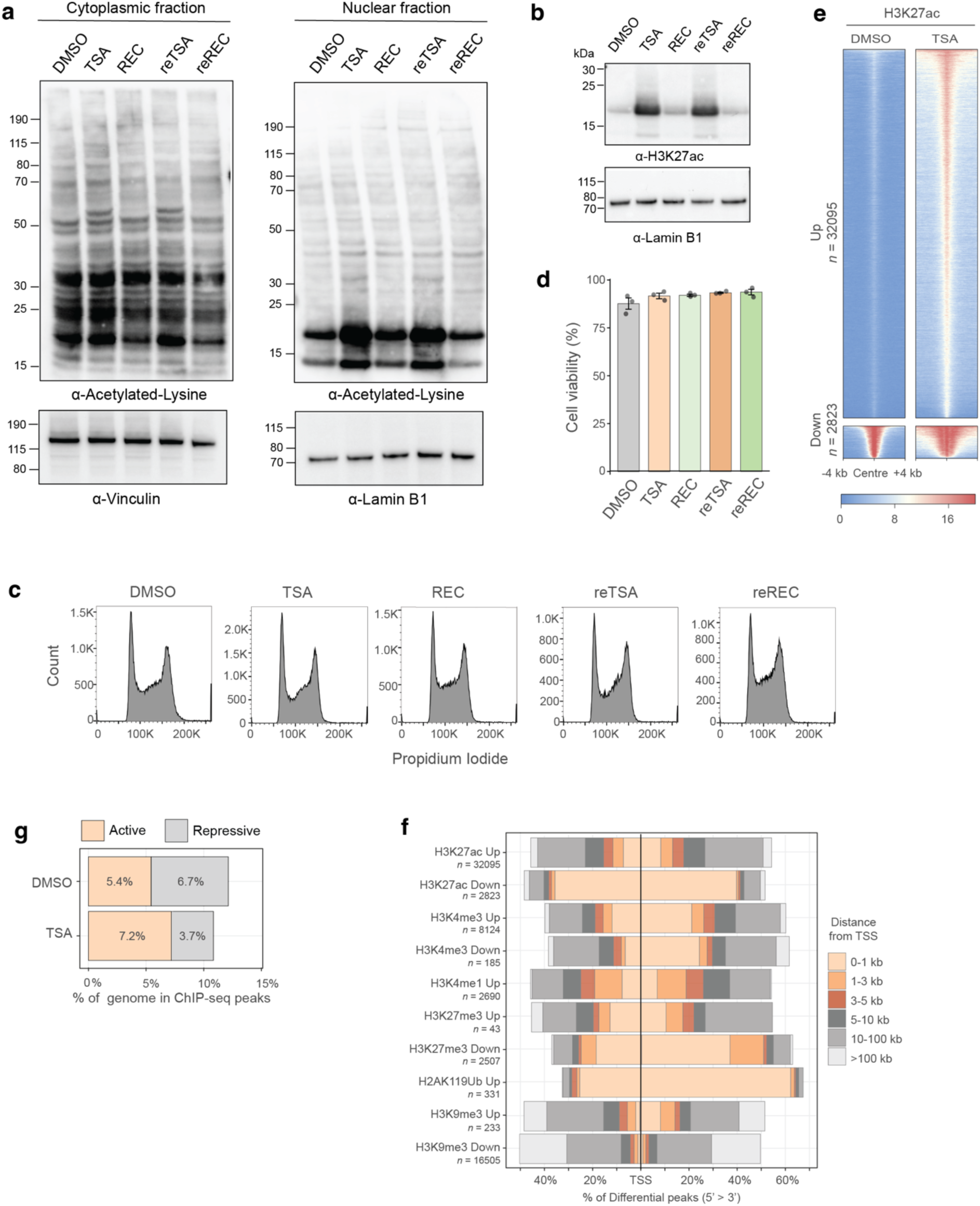
Characterisation of the effect of TSA treatment on H3K27ac levels and histone modification landscape in ESCs. **a**, Western blots showing the levels of lysine acetylation in cytoplasmic (left panels) and nuclear (right panels) extracts in the following conditions: DMSO control, TSA treatment, 24-hour recovery following TSA washout (REC), sequential TSA treatment (reTSA) and 24-hour recovery from the second TSA treatment (reREC). Vinculin and Lamin B1 are used as loading control for cytoplasmic and nuclear samples respectively. **b**, Western blot showing the levels of H3K27 acetylation in nuclear extracts in conditions as in **a**. Lamin B1 is used as loading control. **c**, Cell cycle analysis by flow cytometry using propidium iodode in conditions as in **a**. **d**, Cell cycle viability counts in conditions as in **a**. Mean of three biological replicates ± s.e.m **e**, Heat maps showing H3K27ac ChIP-seq signal at differential peaks. **g**, Bar plots showing percentage of the genome in ChIP-seq peaks of active and repressive histone marks. **f**, Bar plots showing the distance of differential ChIP-seq peaks (H3K27ac, H3K4me3, H3K4me1, H3K27me3, H2AK119Ub, H3K9me3) from transcription start sites (TSS).

**Extended Data Figure 2.**
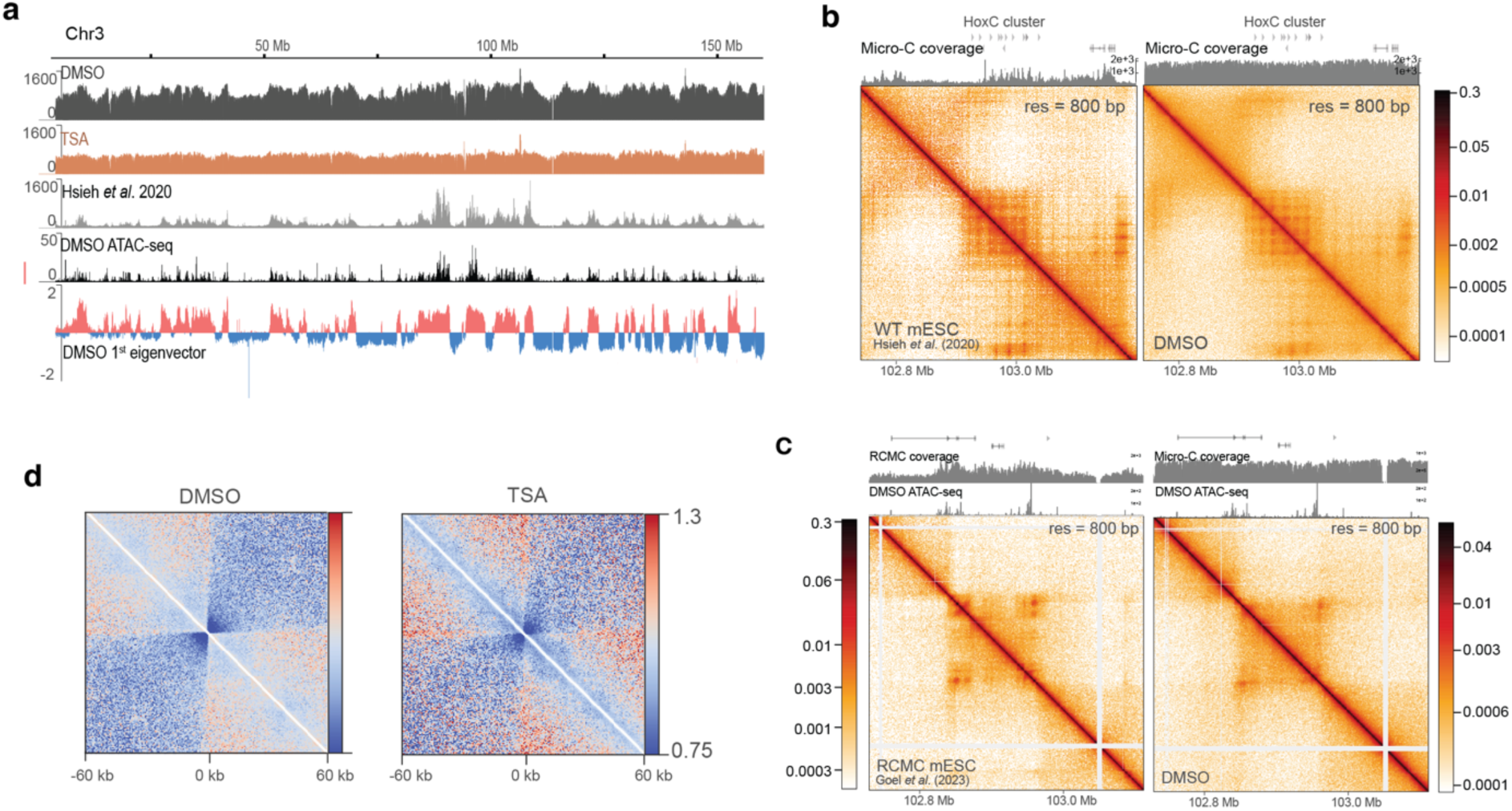
Micro-C data coverage. **a**, Total Micro-C coverage in DMSO, TSA, data from Hsieh *et al*. (2020) on chromosome 3, along with ATAC-seq signal and eigenvector tracks in DMSO. **b**, Micro-C maps showing contacts over the HoxC cluster in WT mESCs (Hsieh *et al*. 2020) and in DMSO. Corresponding Micro-C coverage tracks are displayed above the maps. **c**, Region Capture Micro-C (RCMC) map (left) and Micro-C map (right) from this study showing contacts over the HoxC cluster in WT mESCs (Goel *et al*. 2023) and in DMSO. RCMC data has been downsampled to match depth of the Micro-C map in the capture region. Corresponding RCMC or Micro-C coverage tracks are displayed above the maps. **d**, On-diagonal pile-ups centred at microcompartment loop anchors identified by Goel *et al*. (2023) at the Klf1, Sox2 and Nanog loci (resolution = 600 bp).

**Extended Data Figure 3.**
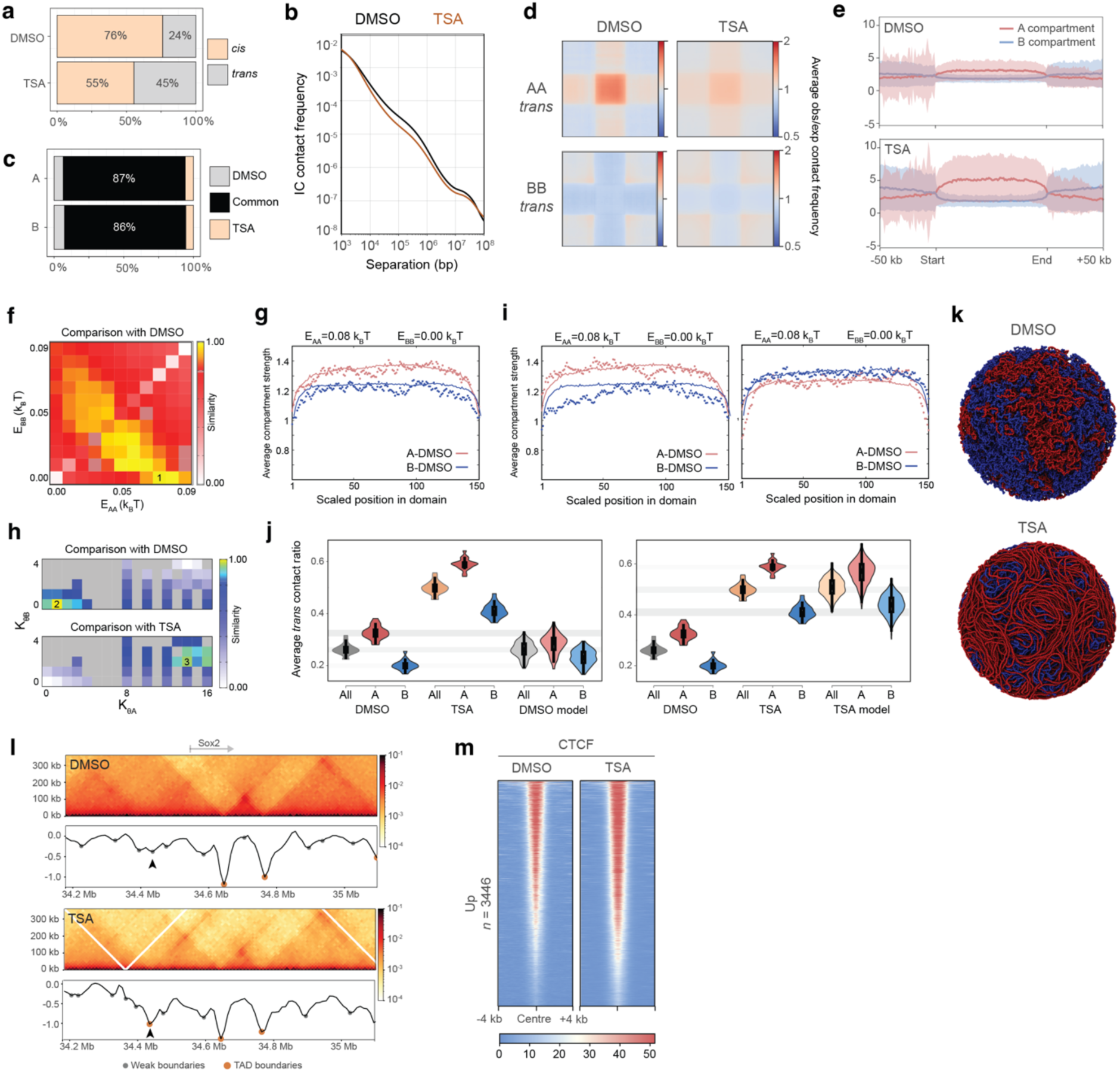
The effect of HDAC inhibition on genome folding. **a**, Ratio of *cis* versus *trans* contacts in Micro-C datasets. **b**, Micro-C contact frequency plotted against genomic separation representing *cis* decay. **c**, Proportion of unique versus common genomic regions assigned to A and B compartments in DMSO and TSA. **d**, Aggregate plots of homotypic interactions between and A and B compartments in *trans*. **e**, Metaplots showing H3K4me1 ChIP-seq read density over A and B compartments (bin size = 1 kb). Shading represents standard deviation. **f**, Heatmap of the similarity scores between the compartment-strength (CS) profiles in DMSO Micro-C and the models for different parameter sets (E_AA_,E_BB_). **g**, A- and B-specific CS profiles from DMSO Micro-C data (lines) and single-chain simulations with optimised attraction energies (points). **h**, Heatmap of the similarity scores between the median of the chromosome-averaged *trans*-contact ratio (TR) in DMSO and TSA Micro-C datasets and the models for different parameter sets (K_θA_,K_θB_). The compartment-specific attraction energies (E_AA_,E_BB_) were maintained equal to the single-chain optimised values. Gray entries indicate untested parameter sets. **i**, Distribution of the *trans*-contact ratios per chromosome in DMSO and TSA Micro-C data, and the optimal models for DMSO (left) and TSA (right). **j**, Distribution of the TR in optimized models for DMSO (left) and TSA (right). No further optimisation of the compartment-specific attractions (E_AA_,E_BB_) was performed to generate the model profiles in TSA (right). Shading represents inter-quartile range. **k**, Example configurations of modelled nuclei corresponding to DMSO (top) and TSA (bottom) conditions. Red beads represent A compartments, blue beads represent B compartments. **l,** Micro-C maps and insulation curves showing a new TAD boundary forming at the Sox2 locus upon TSA treatment (resolution = 10 kb). **m**, Heat maps showing CTCF ChIP-seq signal at differential peaks.

**Extended Data Figure 4.**
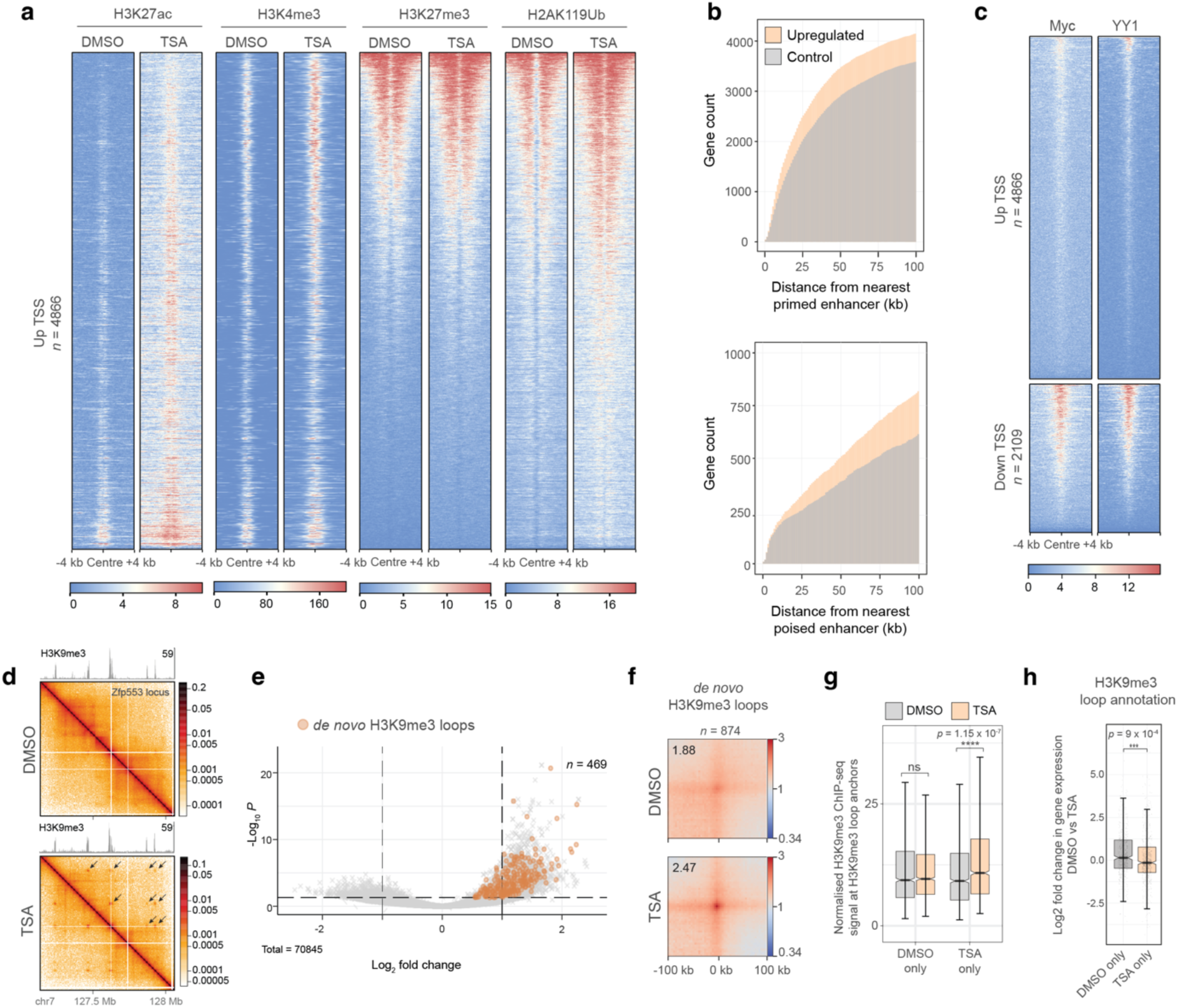
Chromatin changes in TSA near deregulated TSSs. **a**, Heatmaps showing H3K27ac, H3K4me3, H3K27me3 and H2AK119Ub signal in DMSO and TSA around transcription start sites (TSS) of upregulated genes. **b**, Cumulative histogram showing genomic distance between upregulated gene promoters and the nearest primed (upper panel) or poised (lower panel) enhancer. Control genes represent an expression-matched gene set that does not increase in expression. **c**, Heatmaps showing Myc and YY1 ChIP-seq signal around up- and downregulated TSSs. **d**, Contact map showing extensive H3K9me3-associated differential looping at the Zfp553 locus. Corresponding H3K9me3 signal is displayed above. **e**, Volcano plot of differential loops between DMSO and TSA with *de novo* H3K9me3 TSA loops highlighted in orange. Positive log_2_ fold change indicates stronger interaction in TSA. **f**, Pile-up of Micro-C signal (resolution = 4 kb) around *de novo* H3K9me3 loops that form in TSA. **g**, Boxplot showing the normalised H3K9me3 signal at anchors of *de novo* H3K9me3 TSA loops. Data shown are the median, with hinges corresponding to interquartile range (IQR) and whiskers extending to the lowest and highest values within 1.5× IQR, notch shows 95% confidence interval (CI) (uppaired two-tailed t-test; ns > 0.05, ****p < 0.0001). **h**, Log_2_ fold change in gene expression (TSA vs DMSO) of genes that are nearest DMSO-only or TSA-only loop anchors. Data shown are the median, with hinges corresponding to IQR and whiskers extending to the lowest and highest values within 1.5× IQR, notch shows 95% CI (unpaired two-tailed t-test; ns > 0.05, **p < 0.01).

**Extended Data Figure 5.**
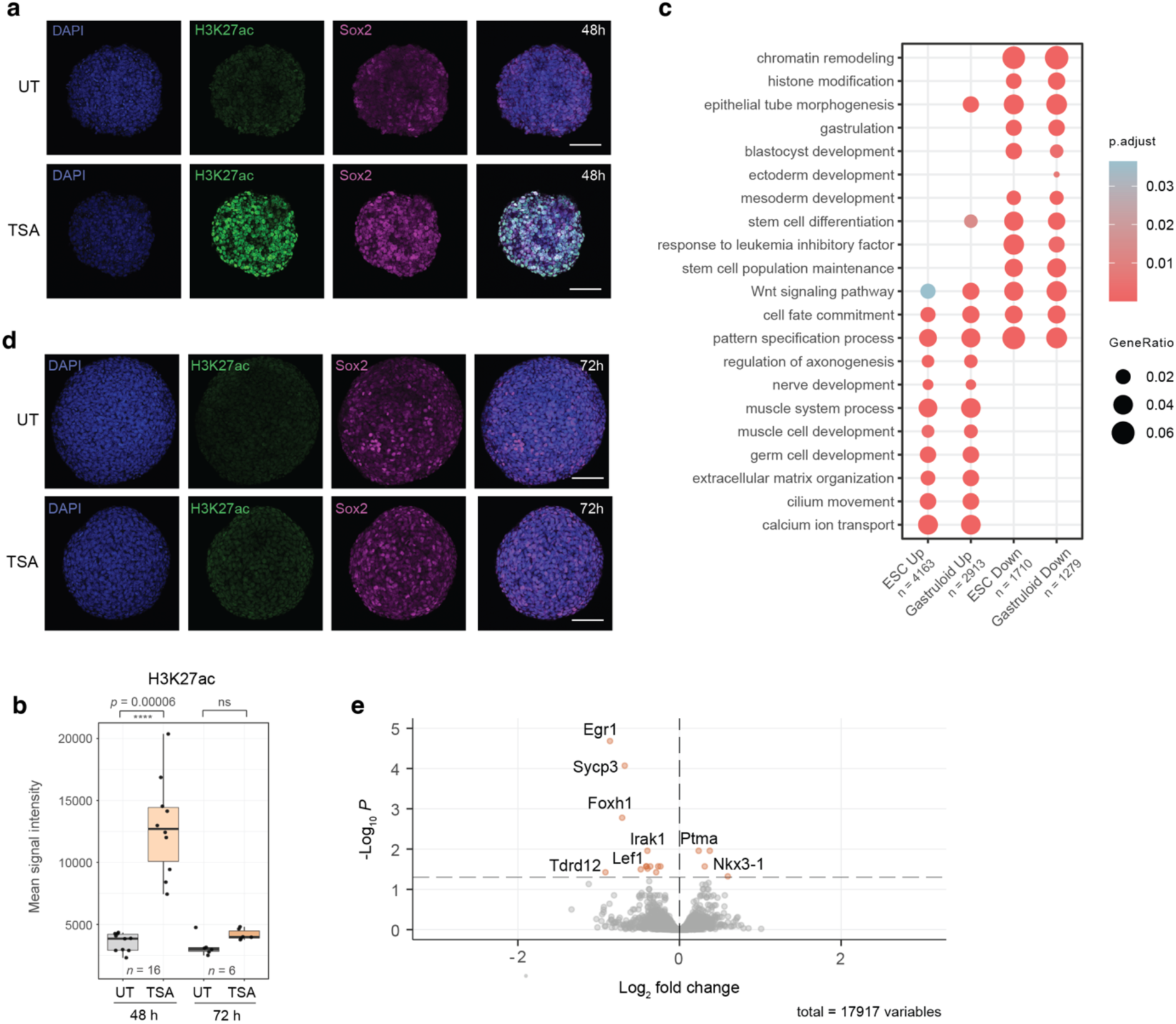
HDAC inhibition-induced chromatin and gene expression changes in mESC and gastruloids. **a**, Representative images of H3K27ac and Sox2 immunostaining in 48-hour gastruloids with (bottom) or without (top) TSA treatment (scale bar = 100 *μ*m). **b**, Quantification of H3K27ac (left panel) and Sox2 (right panel) signal intensities in early (48 h, 72 h) gastruloids with and without TSA treatment. Data shown are the median, with hinges corresponding to IQR and whiskers extending to the lowest and highest values within 1.5× IQR. **c**, Gene Ontology enrichment of terms related to development among differentially expressed genes in mESCs and 48-hour gastruloids after 4 hours of TSA treatment. **d**, Representative images of H3K27ac and Sox2 immunostaining in 72-hour gastruloids with (bottom) or without (top) TSA treatment (scale bar = 100 *μ*m). **e**, Volcano plot showing differentially expressed genes in 120-hour gastruloids with versus without TSA treatment (significance cutoff: adjusted *p*-value > 0.05) Positive log_2_ fold change signifies upregulation in TSA-treated gastruloids.

**Extended Data Figure 6.**
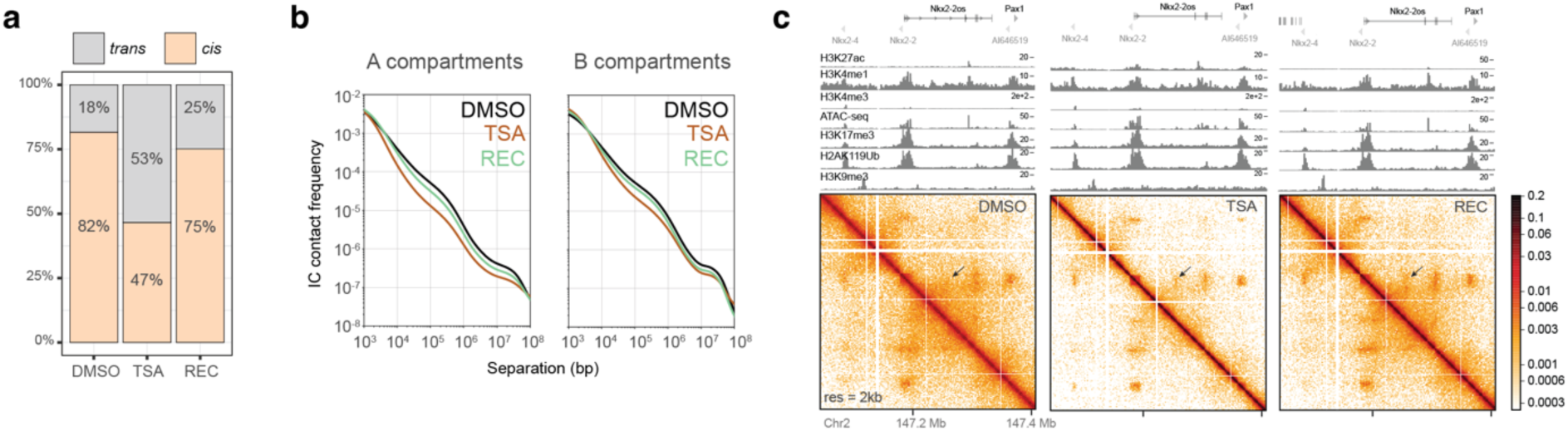
Reversibility of HDAC inhibition-induced architectural changes. **a**, Ratio of *cis* versus *trans* contacts in Micro-C datasets. **b**, Contact frequency in Micro-C plotted against genomic separation representing *cis* decay by compartment. **c**, Micro-C maps at the Nkx2-2 locus showing incomplete architectural recovery. ChIP-seq tracks of the corresponding condition are shown above.

**Extended Data Figure 7.**
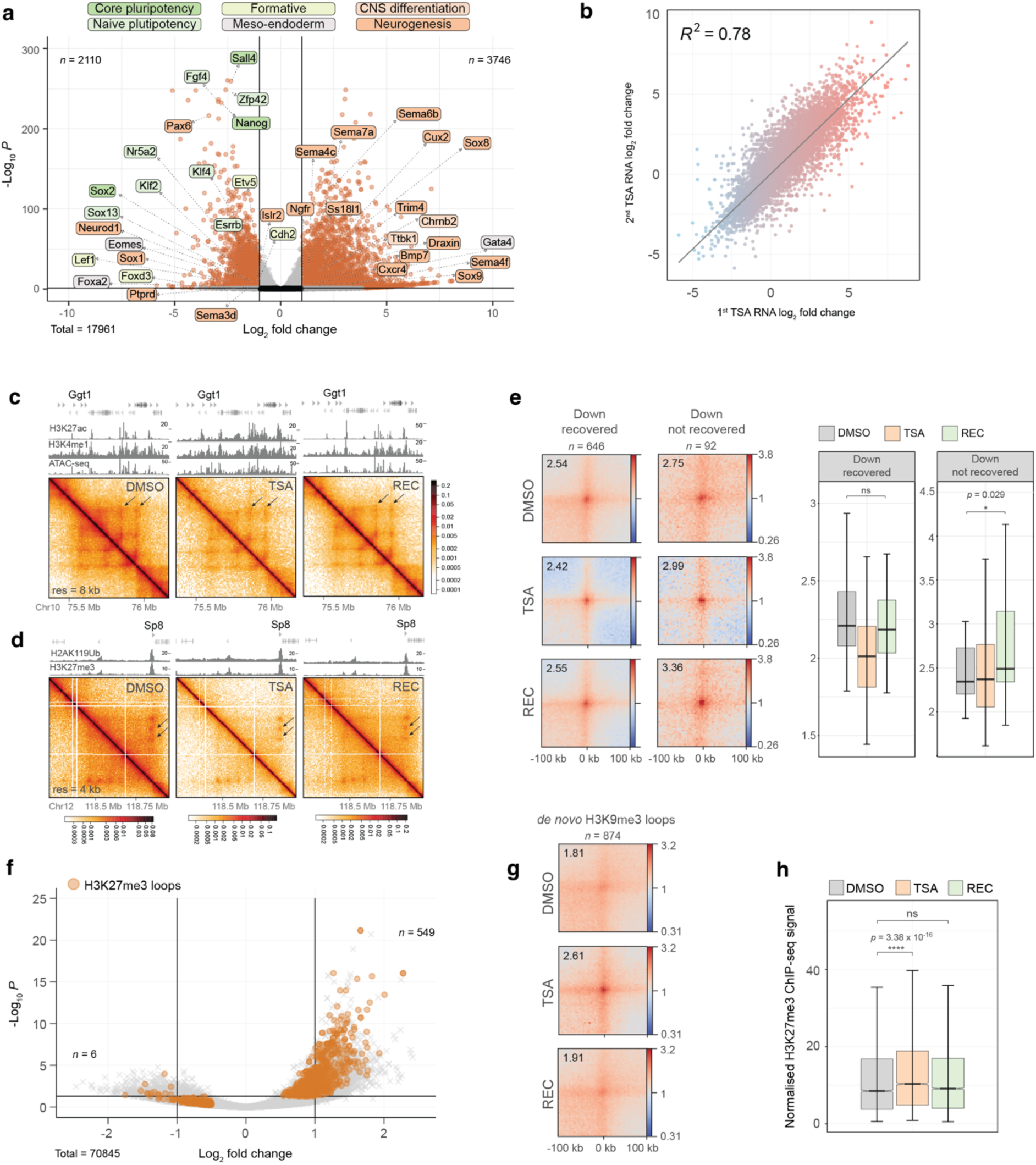
A second TSA treatment triggers memory in mESCs. **a**, Volcano plot showing differential gene expression (significance cutoffs: adjusted *p*-value > 0.05, absolute log_2_ fold change > 1) upon reTSA treatment. Labels correspond to core and naïve pluripotency, formative, meso-endodermal, CNS differentiation and neurogenesis marker genes. Positive log_2_ fold change signifies upregulation in upon reTSA treatment. **b**, Scatterplot showing the correlation of transcriptomic changes induced by the first and second TSA treatments. **c**, Contact map showing enhancer-promoter contacts at the Ggt1 locus that shows sustained transcriptional upregulation. Corresponding H3K27ac, H3K4me1 and ATAC-seq signal are displayed above. **d**, Contact map showing Polycomb-mediated contacts at the Sp8 locus that shows sustained transcriptional downregulation. Corresponding H2AK119Ub and H3K27me3 signal are displayed above. **e**, Aggregate plots of Micro-C signal around loops where anchors overlap with downregulated TSSs that recover (left panes) or do not recover (right panels) (resolution = 4 kb). Quantification of piled-up loop signal is shown on the right (paired two-tailed t-test; ns > 0.05, ***p < 0.001). Data shown are the median, with hinges corresponding to IQR and whiskers extending to the lowest and highest values within 1.5× IQR. **f**, Volcano plot of differential loops between DMSO and TSA. H3K27me3 loops are highlighted in orange. Positive log_2_ fold change indicates stronger interaction in TSA. **g**, Pile-up of Micro-C signal (resolution = 4 kb) around *de novo* H3K9me3 loops that form in TSA. **h**, Boxplot showing the normalised H3K27me3 signal at anchors of H3K27me3 loops in DMSO, TSA and Recovery. Data shown are the median, with hinges corresponding to IQR and whiskers extending to the lowest and highest values within 1.5× IQR (unpaired two-tailed t-test; ns > 0.05, ***p < 0.001).

## Notes

https://github.com/cavallifly/Paldi_et_al_2024

https://www.ncbi.nlm.nih.gov/geo/query/acc.cgi?acc=GSE281151

### Competing Interest Statement

The authors have declared no competing interest.

## References

1. Schlesinger, S. & Meshorer, E. Open Chromatin, Epigenetic Plasticity, and Nuclear Organization in Pluripotency. Dev Cell 48, 135–150 (2019).

2. Pachano, T., Crispatzu, G. & Rada-Iglesias, A. Polycomb proteins as organizers of 3D genome architecture in embryonic stem cells. Brief Funct Genomics 18, 358– 366 (2019).

3. Macrae, T. A., Fothergill-Robinson, J. & Ramalho-Santos, M. Regulation, functions and transmission of bivalent chromatin during mammalian development. Nat Rev Mol Cell Biol 24, 6–26 (2023).

4. Urso, A. D.’ & Brickner, J. H. Mechanisms of epigenetic memory. (2014) doi:10.1016/j.tig.2014.04.004.

5. Atlasi, Y. & Stunnenberg, H. G. The interplay of epigenetic marks during stem cell dìerentiation and development. Nat Rev Genet 18, 643–658 (2017).

6. Yoshida, M., Kijima, M., Akita, M. & Beppu, T. Potent and specific inhibition of mammalian histone deacetylase both in vivo and in vitro by trichostatin A. Journal of Biological Chemistry 265, 17174–17179 (1990).

7. Willemin, A., Szabó, D. & Pombo, A. Epigenetic regulatory layers in the 3D nucleus. Mol Cell 84, 415–428 (2024).

8. Piunti, A. & Shilatifard, A. Epigenetic balance of gene expression by polycomb and compass families. Science (1979) 352, (2016).

9. Lagger, G. et al. Essential function of histone deacetylase 1 in proliferation control and CDK inhibitor repression. EMBO Journal 21, 2672–2681 (2002).

10. Dovey, O. M., Foster, C. T. & Cowley, S. M. Histone deacetylase 1 (HDAC1), but not HDAC2, controls embryonic stem cell dìerentiation. Proc Natl Acad Sci U S A 107, 8242–8247 (2010).

11. Karantzali, E. et al. Histone deacetylase inhibition accelerates the early events of stem cell dìerentiation: Transcriptomic and epigenetic analysis. Genome Biol 9, (2008).

12. Kidder, B. L. & Palmer, S. HDAC1 regulates pluripotency and lineage specific transcriptional networks in embryonic and trophoblast stem cells. Nucleic Acids Res 40, 2925–2939 (2012).

13. Jamaladdin, S. et al. Histone deacetylase (HDAC) 1 and 2 are essential for accurate cell division and the pluripotency of embryonic stem cells. Proc Natl Acad Sci U S A 111, 9840–9845 (2014).

14. Fejes Tóth, K., et al. Trichostatin A-induced histone acetylation causes decondensation of interphase chromatin. J Cell Sci 117, 4277–4287 (2004).

15. Otterstrom, J. et al. Super-resolution microscopy reveals how histone tail acetylation àects DNA compaction within nucleosomes in vivo. Nucleic Acids Res 47, 8470–8484 (2019).

16. Szabo, Q. et al. Regulation of single-cell genome organization into TADs and chromatin nanodomains. Nat Genet 52, 1151–1157 (2020).

17. Hsieh, T.-H. S. et al. Resolving the 3D Landscape of Transcription-Linked Mammalian Chromatin Folding. Mol Cell 1–15 (2020) doi:10.1016/j.molcel.2020.03.002.

18. Hsieh, T. H. S. et al. Enhancer–promoter interactions and transcription are largely maintained upon acute loss of CTCF, cohesin, WAPL or YY1. Nat Genet 54, 1919–1932 (2022).

19. Goel, V. Y., Huseyin, M. K. & Hansen, A. S. Region Capture Micro-C reveals coalescence of enhancers and promoters into nested microcompartments. 1–8 (2022).

20. Bonev, B. et al. Multiscale 3D Genome Rewiring during Mouse Neural Development. Cell 171, 557–572.e24 (2017).

21. Leidescher, S. et al. Spatial organization of transcribed eukaryotic genes. Nat Cell Biol 24, 327–339 (2022).

22. Orsi, G. A. et al. Biophysical ordering transitions underlie genome 3D re-organization during cricket spermiogenesis. Nat Commun 14, 1–16 (2023).

23. Di Stefano, M. & Cavalli, G. Integrative studies of 3D genome organization and chromatin structure. Curr Opin Struct Biol 77, 102493 (2022).

24. Girard, M., De La Cruz, M. O., Marko, J. F. & Erbaş, A. Heterogeneous flexibility can contribute to chromatin segregation in the cell nucleus. Phys Rev E 110, 1–12 (2024).

25. Cruz-Molina, S. et al. PRC2 Facilitates the Regulatory Topology Required for Poised Enhancer Function during Pluripotent Stem Cell Dìerentiation. Cell Stem Cell 20, 689–705.e9 (2017).

26. Crispatzu, G. et al. The chromatin, topological and regulatory properties of pluripotency-associated poised enhancers are conserved in vivo. Nat Commun 12, 1–17 (2021).

27. Fraser, J. et al. Hierarchical folding and reorganization of chromosomes are linked to transcriptional changes in cellular dìerentiation. Mol Syst Biol 11, 1–14 (2015).

28. Winick-Ng, W. et al. Cell-type specialization is encoded by specific chromatin topologies. Nature 599, 684–691 (2021).

29. Yao, Y.-L., Yang, W.-M. & Seto, E. Regulation of Transcription Factor YY1 by Acetylation and Deacetylation. Mol Cell Biol 21, 5979–5991 (2001).

30. Faiola, F. et al. Dual Regulation of c-Myc by p300 via Acetylation-Dependent Control of Myc Protein Turnover and Coactivation of Myc-Induced Transcription. Mol Cell Biol 25, 10220–10234 (2005).

31. Van Den Brink, S. C., et al. Symmetry breaking, germ layer specification and axial organisation in aggregates of mouse embryonic stem cells. Development (Cambridge) 141, 4231–4242 (2014).

32. Beccari, L. et al. Multi-axial self-organization properties of mouse embryonic stem cells into gastruloids. Nature 562, 272–276 (2018).

33. Harris, H. L. et al. Chromatin alternates between A and B compartments at kilobase scale for subgenic organization. Nat Commun 14, (2023).

34. Argelaguet, R. et al. Multi-omics profiling of mouse gastrulation at single-cell resolution. Nature 576, 487–491 (2019).

35. Zhang, Y. et al. Dynamic epigenomic landscapes during early lineage specification in mouse embryos. Nat Genet 50, 96–105 (2018).

36. Xie, W. et al. Epigenomic analysis of multilineage dìerentiation of human embryonic stem cells. Cell 153, 1134–1148 (2013).

37. Gìord, C. A., et al. Transcriptional and epigenetic dynamics during specification of human embryonic stem cells. Cell 153, 1149–1163 (2013).

38. Creyghton, M. P. et al. Histone H3K27ac separates active from poised enhancers and predicts developmental state. Proc Natl Acad Sci U S A 107, 21931–21936 (2010).

39. Rada-Iglesias, A. et al. A unique chromatin signature uncovers early developmental enhancers in humans. Nature 470, 279–285 (2011).

40. Kim, H. S. et al. Pluripotency factors functionally premark cell-type-restricted enhancers in ES cells. Nature 556, 510–514 (2018).

41. Schick, S. et al. Acute BAF perturbation causes immediate changes in chromatin accessibility. Nat Genet 53, 269–278 (2021).

42. Iurlaro, M. et al. Mammalian SWI/SNF continuously restores local accessibility to chromatin. Nat Genet 53, 279–287 (2021).

43. Ghavi-Helm, Y. et al. Enhancer loops appear stable during development and are associated with paused polymerase. Nature 512, 96–100 (2014).

44. Benabdallah, N. S. et al. Decreased Enhancer-Promoter Proximity Accompanying Enhancer Activation. Mol Cell 76, 473–484.e7 (2019).

45. Ing-Simmons, E. et al. Independence of chromatin conformation and gene regulation during Drosophila dorsoventral patterning. Nat Genet 53, 487–499 (2021).

46. Pollex, T. et al. Enhancer–promoter interactions become more instructive in the transition from cell-fate specification to tissue dìerentiation. Nat Genet 56, 686– 696 (2024).

47. Jin, F. et al. A high-resolution map of the three-dimensional chromatin interactome in human cells. Nature 503, 290–294 (2013).

48. Schoenfelder, S. et al. Polycomb repressive complex PRC1 spatially constrains the mouse embryonic stem cell genome. Nat Genet 47, 1179–1186 (2015).

49. Stadhouders, R. et al. Transcription factors orchestrate dynamic interplay between genome topology and gene regulation during cell reprogramming. Nat Genet 50, 238–249 (2018).

50. Paliou, C. et al. Preformed chromatin topology assists transcriptional robustness of Shh during limb development. Proc Natl Acad Sci U S A 116, 12390–12399 (2019).

51. Holoch, D., et al. A Cis-Acting Mechanism Mediates Transcriptional Memory at Polycomb Target Genes in Mammals. Nature Genetics vol. 53 (Springer US, 2021).

52. Margueron, R. et al. Ezh1 and Ezh2 Maintain Repressive Chromatin through Dierent Mechanisms. Mol Cell 32, 503–518 (2008).

53. Lau, M. S. et al. Mutation of a nucleosome compaction region disrupts Polycomb-mediated axial patterning. Science (1979) 355, 1081–1084 (2017).

54. Lanzuolo, C., Roure, V., Dekker, J., Bantignies, F. & Orlando, V. Polycomb response elements mediate the formation of chromosome higher-order structures in the bithorax complex. Nat Cell Biol 9, 1167–1174 (2007).

55. Bantignies, F. et al. Polycomb-dependent regulatory contacts between distant hox loci in drosophila. Cell 144, 214–226 (2011).

56. Cheutin, T. & Cavalli, G. Loss of PRC1 induces higher-order opening of Hox loci independently of transcription during Drosophila embryogenesis. Nat Commun 9, 1–11 (2018).

57. Ogiyama, Y., Schuettengruber, B., Papadopoulos, G. L., Chang, J. M. & Cavalli, G. Polycomb-Dependent Chromatin Looping Contributes to Gene Silencing during Drosophila Development. Mol Cell 71, 73–88.e5 (2018).

58. Mas, G. et al. Promoter bivalency favors an open chromatin architecture in embryonic stem cells. Nat Genet 50, 1452–1462 (2018).

59. Murphy, S. E. & Boettiger, A. N. Polycomb repression of Hox genes involves spatial feedback but not domain compaction or phase transition. Nat Genet 56, 493–504 (2024).

60. Bártová, E. et al. Nuclear levels and patterns of histone H3 modification and HP1 proteins after inhibition of histone deacetylases. J Cell Sci 118, 5035–5046 (2005).

61. Mahy, N. L., Perry, P. E. & Bickmore, W. A. Gene density and transcription influence the localization of chromatin outside of chromosome territories detectable by FISH. Journal of Cell Biology 159, 753–763 (2002).

62. Chambeyron, S., Da Silva, N. R., Lawson, K. A. & Bickmore, W. A. Nuclear re-organisation of the Hoxb complex during mouse embryonic development. Development 132, 2215–2223 (2005).

63. Brown, J. M. et al. Association between active genes occurs at nuclear speckles and is modulated by chromatin environment. Journal of Cell Biology 182, 1083– 1097 (2008).

64. Hildebrand, E. M. & Dekker, J. Mechanisms and Functions of Chromosome Compartmentalization. Trends Biochem Sci 45, 385–396 (2020).

65. Xiao, J., Hafner, A. & Boettiger, A. N. How subtle changes in 3d structure can create large changes in transcription. Elife 10, 1–27 (2021).

66. Di Stefano, M., Nützmann, H. W., Marti-Renom, M. A. & Jost, D. Polymer modelling unveils the roles of heterochromatin and nucleolar organizing regions in shaping 3D genome organization in Arabidopsis thaliana. Nucleic Acids Res 49, 1840–1858 (2021).

67. Hoencamp, C. et al. 3D genomics across the tree of life reveals condensin II as a determinant of architecture type. Science (1979) 372, 984–989 (2021).

68. Owen, J. A., Osmanović, D. & Mirny, L. Design principles of 3D epigenetic memory systems. Science (1979) 382, (2023).

69. Jost, D. & Vaillant, C. Epigenomics in 3D: Importance of long-range spreading and specific interactions in epigenomic maintenance. Nucleic Acids Res 46, 2252– 2264 (2018).

70. Michieletto, D., Orlandini, E. & Marenduzzo, D. Polymer model with epigenetic recoloring reveals a pathway for the de novo establishment and 3D organization of chromatin domains. Phys Rev X 6, 1–15 (2016).

## Method References

71. Beccari, L. et al. Generating Gastruloids from Mouse Embryonic Stem Cells. Protoc Exch 1–8 (2018) doi:10.1038/protex.2018.094.

72. Vianello, S., Girgin, M., Rossi, G. & Lutolf, M. Protocol to immunostain Gastruloids (LSCB, EPFL). Protocols.Io 1–10 (2020).

73. Andrey, G. et al. Characterization of hundreds of regulatory landscapes in developing limbs reveals two regimes of chromatin folding. Genome Res 27, 223–233 (2017).

74. Lee, T. I., Johnstone, S. E. & Young, R. A. Chromatin immunoprecipitation and microarray-based analysis of protein location. 1, 729–748 (2006).

75. Radzisheuskaya, A. et al. Complex-dependent histone acetyltransferase activity of KAT8 determines its role in transcription and cellular homeostasis. Mol Cell 81, 1749–1765.e8 (2021).

76. Love, M. I., Huber, W. & Anders, S. Moderated estimation of fold change and dispersion for RNA-seq data with DESeq2. Genome Biol 15, 1–21 (2014).

77. Yu, G., Wang, L. G., Han, Y. & He, Q. Y. ClusterProfiler: An R package for comparing biological themes among gene clusters. OMICS 16, 284–287 (2012).

78. Langmead, B. & Salzberg, S. L. Fast gapped-read alignment with Bowtie 2. Nat Methods 9, 357–359 (2012).

79. Li, H. et al. The Sequence Alignment/Map format and SAMtools. Bioinformatics 25, 2078–2079 (2009).

80. Tarasov, A., Vilella, A. J., Cuppen, E., Nijman, I. J. & Prins, P. Sambamba: Fast processing of NGS alignment formats. Bioinformatics 31, 2032–2034 (2015).

81. Fursova, N. A. et al. Synergy between Variant PRC1 Complexes Defines Polycomb-Mediated Gene Repression. Mol Cell 74, 1020–1036.e8 (2019).

82. Ramírez, F., Dündar, F., Diehl, S., Grüning, B. A. & Manke, T. DeepTools: A flexible platform for exploring deep-sequencing data. Nucleic Acids Res 42, 187–191 (2014).

83. Robinson, J. T., Thorvaldsdottir, H., Turner, D. & Mesirov, J. P. igv.js: an embeddable JavaScript implementation of the Integrative Genomics Viewer (IGV). Bioinformatics 39, 23–24 (2023).

84. Kerpedjiev, P. et al. HiGlass: Web-based visual exploration and analysis of genome interaction maps. Genome Biol 19, (2018).

85. Zhang, Y. et al. Model-based Analysis of ChIP-Seq (MACS). Genome Biol 9, R137 (2008).

86. Ross-Innes, C. S. et al. Dìerential oestrogen receptor binding is associated with clinical outcome in breast cancer. Nature 481, 389–393 (2012).

87. Yu, G., Wang, L. G. & He, Q. Y. ChIP seeker: An R/Bioconductor package for ChIP peak annotation, comparison and visualization. Bioinformatics 31, 2382–2383 (2015).

88. Chronis, C. et al. Cooperative Binding of Transcription Factors Orchestrates Reprogramming. Cell 168, 442–459.e20 (2017).

89. Servant, N. et al. HiC-Pro: An optimized and flexible pipeline for Hi-C data processing. Genome Biol 16, (2015).

90. Abdennur, N. & Mirny, L. A. Cooler: Scalable storage for Hi-C data and other genomically labeled arrays. Bioinformatics 36, 311–316 (2020).

91. Yang, T. et al. HiCRep: assessing the reproducibility of Hi-C data using a stratum-adjusted correlation coèicient. Genome Res 27, 1939–1949 (2017).

92. Imakaev, M. et al. Iterative correction of Hi-C data reveals hallmarks of chromosome organization. Nat Methods 9, 999–1003 (2012).

93. Roayaei Ardakany, A., Gezer, H. T., Lonardi, S. & Ay, F. Mustache: Multi-scale detection of chromatin loops from Hi-C and Micro-C maps using scale-space representation. Genome Biol 21, 1–17 (2020).

94. Flyamer, I. M., Illingworth, R. S. & Bickmore, W. A. Coolpup.py: Versatile pile-up analysis of Hi-C data. Bioinformatics 36, 2980–2985 (2020).

95. Quinlan, A. R. & Hall, I. M. BEDTools: A flexible suite of utilities for comparing genomic features. Bioinformatics 26, 841–842 (2010).

96. Kremer, K. & Grest, G. S. Dynamics of entangled linear polymer melts: A molecular-dynamics simulation. J Chem Phys 92, 5057–5086 (1990).

97. Di Stefano, M., Paulsen, J., Lien, T. G., Hovig, E. & Micheletti, C. Hi-C-constrained physical models of human chromosomes recover functionally-related properties of genome organization. Sci Rep 6, 1–12 (2016).

98. Plimpton, S. Fast Parallel Algorithms for Short-Range Molecular Dynamics. J Comput Phys 117, 1–19 (1995).

99. Ghosh, S. K. & Jost, D. How epigenome drives chromatin folding and dynamics, insights from èicient coarse-grained models of chromosomes. PLoS Comput Biol 14, 1–26 (2018).

100. Gu, B. et al. Transcription-coupled changes in nuclear mobility of mammalian cis-regulatory elements. Science(1979) 359, 1050–1055 (2018).

101. Humphrey, W., Dalke, A. & Schulten, K. VMD: Visual molecular dynamics. J Mol Graph 14, 33–38 (1996).

